# Voice Changes in Aging Birdsong: A Longitudinal Study in Adult Male Zebra Finches

**DOI:** 10.1101/2025.10.27.684935

**Authors:** Michelle L. Gordon, Arpita Gulati, Robin A. Samlan, Julie E. Miller

## Abstract

By 2030, 1.4 billion people worldwide will be 60 years or older. A significant proportion will experience changes in voice, which can indicate early signs of neurodegenerative disease. Most research and treatment studies have focused on the larynx because of its accessibility and known role in moderating pitch, loudness, and voice quality. Pre-clinical animal models offer important insight into brain mechanisms involved in aging vocalizations. Previously, we conducted a cross-sectional analysis in adult male zebra finches, discovering that song changes occur at transitional timepoints, between young adult, middle age, and older ages. Song changes are accompanied by re-organization of gene networks in a song-dedicated basal ganglia brain nucleus at middle age that degrade in old age. To more precisely examine how aging alters songs during these transitional timepoints, we followed individual male zebra finches longitudinally over adulthood. Measurements of aging human voice, including duration, intensity, smoothed cepstral peak prominence, and fundamental frequency (f_o_ mean, median, standard deviation, and coefficient of variation), were carried out on birdsong syllables. Our results show that changes in f_o_ mean and intensity are the measurements most sensitive to aging and consistent across multiple mornings in the finches. Notably, acoustic changes were also more pronounced in the transition from young adult to middle age, a time at which changes in aging voice also emerge. Our findings strengthen the validity of the finch model for use in aging research and motivate future investigations exploring the relative contributions of laryngeal and brain mechanisms to aging vocalizations.

## Introduction

By 2030, one in six people worldwide will be 60 years of age or older (World Health Organization, 2024). In the United States, this represents 24% of the population, with a doubling projected by 2050 (U.S. Census Bureau, 2023; PRB, 2024). In 2022, 17 states had a median age of 40 or above, pointing to a rapidly growing middle age population, a time when neurological diseases can develop (PRB, 2024). At least 30% of adults older than 50 years of age (Rojas et al., 2020) experience changes in their voice that impact communication, participation in employment, and quality of life (Beton et al., 2023; Etter et al., 2013; Lindström et al., 2023; Wang et al., 2023). Presbyphonia, or age-related vocal disorders, include a quiet, breathy, rough voice, shortened phrases, and sex-dependent changes in pitch (Cavallaro et al., 2024; Taylor et al., 2020). Several changes in vocal fold vibration can occur with aging, such as incomplete glottal closure; altered open phase or amplitude of vibration; and decreased periodicity or symmetry (Ahmad et al., 2012; Biever & Bless, 1989; Kosztyła-Hojna et al., 2023; Samlan et al., 2016; Samlan et al., 2018; Stager et al., 2021; Yamauchi et al., 2012; Yamauchi et al., 2015); the brain’s role has not been explored.

Measurements of voice are often used to describe aging voice. They include mean intensity (a correlate of loudness). Correlates of voice quality include cepstral peak prominence (periodicity), shimmer (cycle to cycle changes in the amplitude of the sound wave), jitter (cycle to cycle frequency changes) and harmonics to noise ratio. Cepstral peak prominence (CPP) and its smoothed version (CPPS) are the most reliable and valid measures of voice quality currently used. A low CPP/CPPS indicates decreased voice quality in the form of breathiness or roughness (Fraile, 2014; Hillenbrand et al., 1994; Hillenbrand & Houde, 1996; Samlan et al., 2013). It has been used to diagnose the presence or absence of a voice disorder with at least 82% accuracy (Brockmann-Bauser et al., 2021; Schultz et al., 2023). The rate of vocal fold vibration and its variability is measured through mean fundamental frequency (f_o_) and standard deviation (std), respectively, and perceived as pitch which becomes monotonous in Parkinson’s Disease, a disease of aging (Bunton, 2006; Harel et al., 2004; Rusz et al., 2021).

While an abundance of literature strongly supports voice and speech changes in aging adults, there is considerable variability in subject age ranges, background (such as vocal training), health and medication status, and variation in measurements used across studies leading to inconsistent findings across studies. Also, cross-sectional population studies are typically the norm. Few studies measure voice and speech changes in the same subject longitudinally over the entire adult lifespan; while vocal quality deteriorates with age, the low number of available subjects along with day to day voice changes are limitations of this research (Frankenberg et al., 2021; Hunter et al., 2012; Tucker et al., 2023; Verdonck-de Leeuw & Mahieu, 2004). Findings are also sex-dependent (Stathopoulos et al., 2011). Despite these multiple confounding factors in studying aging vocalizations, they represent a potentially powerful tool for early detection and diagnosis in a variety of age-related disorders and diseases.

As a starting point, pre-clinical animal models of aging voice offer advantages over human subjects. Subject history and environmental settings can be controlled. The underlying brain mechanisms can be investigated at the molecular, cellular, and circuit levels and related to the vocal output. The adult male zebra finch songbird is an excellent model system for these purposes due to its well-characterized song-dedicated brain circuitry that is absent in females and its face validity to human brain centers, as established by its genetic, anatomical, and physiological similarities (Pfenning et al., 2014; Sakata et al., 2020). Brain pathways for vocal learning and production of birdsong synapse on cranial motor neurons and then project to the syrinx, which is anatomically and functionally similar to the human larynx (Riede & Goller, 2010a,b).

In our prior publications, we made acoustic measurements of intensity, duration, CPPS, and f_o_ in song syllables from a cross-sectional group of adult male finches in our self-sustaining laboratory colony. Age categories were assigned as follows: young adults (∼8 months of age post-hatch), middle age (∼24 months), and older (∼40 months) (Badwal et al., 2020; Badwal et al., 2018; Higgins et al., 2025). Finches are reared and housed in the same room which is under rigorous temperature and lighting controls, and they all receive identical food and supplements. Age at birth is recorded. Health status is monitored throughout their life. Thus, our regulation of environmental factors, diet, and tracking finch health from birth through end of life minimizes the impact of these factors on experimental investigations.

Birdsong consists of a repeated sequence of syllables that form a motif unit (Badwal et al., 2018). Syllables have vowel and consonant-like properties (**Figure 1**). At the syllable level, we discovered that middle age was a key transitional timepoint in birdsong for two cross-sectional studies in adult male zebra finches (Badwal et al., 2020; Higgins et al., 2025). A drop in CPPS scores was detected at middle age compared to younger adults for two syllable types across two studies. Vocal intensity (loudness) also increased with age across syllable types. In one study, we detected reduced syllable duration and f_o_ std in older adults compared to younger adults implying a faster, more monotonous song in older finches (Higgins et al., 2025). We also paired age-related changes in song with mRNA level brain expression patterns in song-dedicated basal ganglia nucleus Area X. Our findings showed that young adults have a tightly organized network of song-associated genes that re-organizes at middle age and noticeably degrades at older ages (Higgins et al., 2025). Despite alterations in the gene network that includes ortholog genes to human disorders (speech, neurodegenerative diseases), the overall structure of the song is preserved. Through acoustic quantification, we detected only subtle changes at the syllable level with aging that could not be visually detected nor apparent to human hearing, suggesting that preservation of song with aging involves compensatory mechanisms.

**Fig 1.**
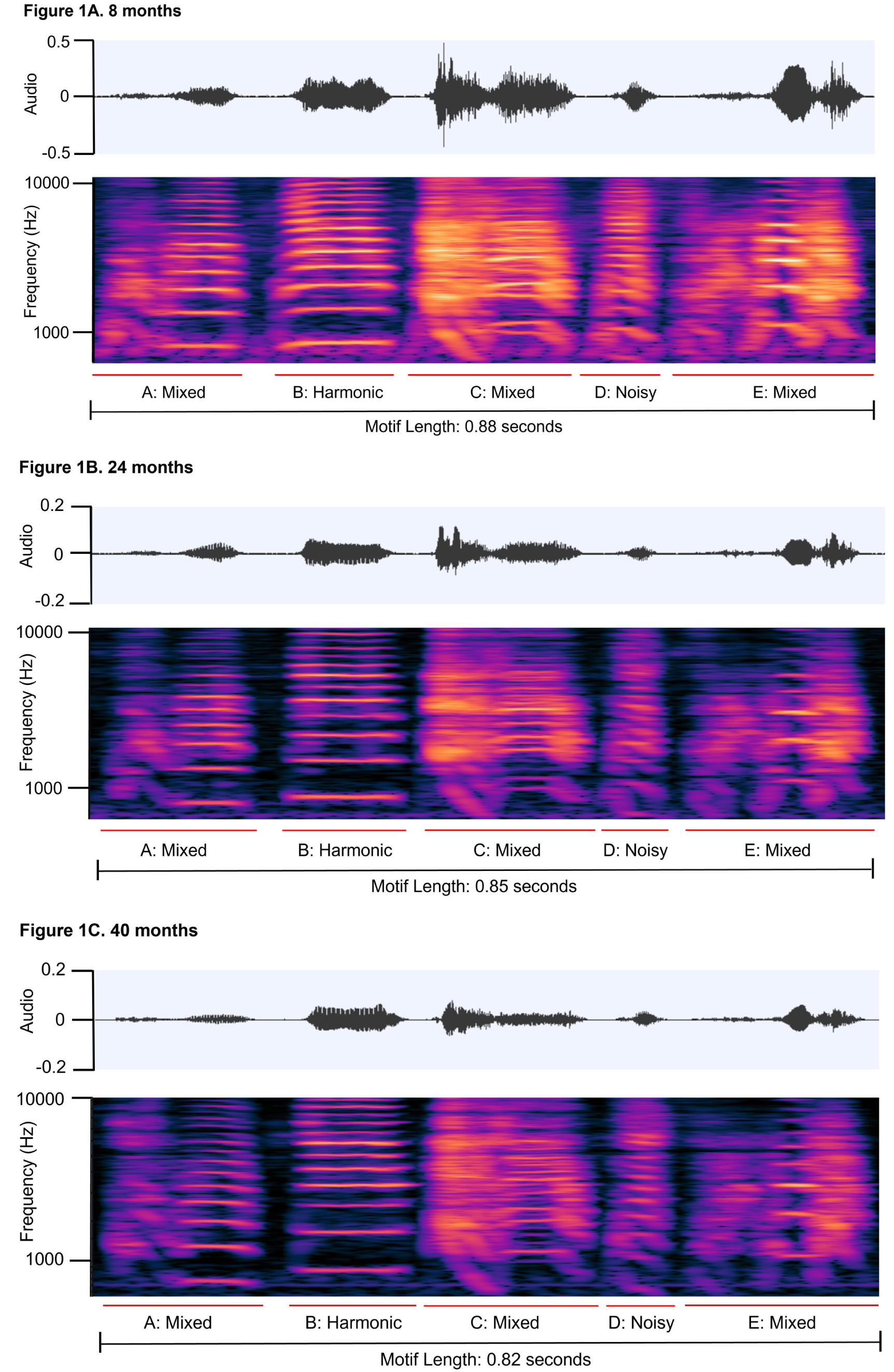
Spectrogram examples of a representative motif from an individual adult male zebra finch (Bird ID: R520) at the three ages. The top row in each panel is the linear amplitude of the audio file while the bottom row is the spectrogram. Frequency (Hertz-Hz) is on the y-axes, and Time (seconds) is on the x-axes. The motif is comprised of syllables labeled A-E and assigned as either harmonic, noisy, or mixed type per our prior categorization (Badwal et al., 2020; Badwal et al., 2018). **A)** Young adult motif at 8 months post-hatch. **B)** Middle age adult motif at 24 months post-hatch. **C)** Older adult motif at 40 months post-hatch. Audio .wav files of these motifs were visualized in the open-source program, Audacity (https://www.audacityteam.org/), and are included in the **Supplemental Material**.

Our current longitudinal study aimed to follow the same group of adult male zebra finches over their four year lifespan to determine if certain acoustic features significantly change within and across finches. We compared the songs of individual finches as they aged from young adults to middle age, and to older ages, using the same age ranges as in our prior studies. We also sampled three consecutive mornings of song at each age category to evaluate day to day variability in acoustic features. Findings were compared to our prior cross-sectional studies and human literature on aging voice.

## Methods

### Subjects and Experimental Timeline

All animal use was approved by the Institutional Care and Use Committee at the University of Arizona. Food and water was provided *ad libitum*. Adult non-breeding male zebra finches (n=6) were selected from a self-sustaining aviary colony at the University of Arizona and grouped together into one cage for the duration of the study spanning the years 2017-2020, except when male finches were individually housed for three consecutive mornings of song recordings. Songs were recorded for three mornings at three different ages, beginning as young adults, eight months of age (corresponding to ∼215-234 days post-hatch-dph), then again at middle age, 24 months (∼685-729 dph), and then at older ages, 40 months (∼1178-1221 dph), using pre-defined age ranges as in our prior cross-sectional studies (Badwal et al., 2020; Higgins et al., 2025).

### Song Acquisition and Recording

Song recording followed the same procedure as in Badwal et al (2020) and are summarized here. Adult male zebra finches were raised in an aviary colony under 14 hours of light and 10 hours of darkness and then acclimated to sound attenuation chambers for two days prior to the recording. Recordings were from individual male finches practicing their song in the absence of a female (known as undirected song). All songs were recorded for two hours after lights-on in the morning. Songs were acquired using free-field omnidirectional microphones (Shure 93) suspended at set points over the center of the cage, connected to an audiobox (Audiobox: 44.1 kHz sampling rate per 24-bit depth; Niles, IL) and recorded with Sound Analysis Pro 2011 (SAP 2011, (Tchernichovski et al., 2000) http://soundanalysispro.com/). For each sound chamber, we adhered to our prior methods, in which intensity levels were calculated by playing a 1,000-Hz tone within each chamber and using a type II sound level meter to derive calibration factors for each chamber applied to the raw intensity measurements (Badwal et al., 2020; Badwal et al., 2018; in **Supplemental Tables 2-7**).

### Song Visualization and Extraction

Songs of adult male finches are comprised of repeated stereotyped sequences, motifs, separated by silent periods. Within each motif, there are individual sound known as syllables that have different characteristics (Lachlan et al., 2016). Motifs and syllables (5-6 syllables) were identified and exported from the open source program Praat (https://www.fon.hum.uva.nl/praat/) (Boersma & Van Heuven, 2001). The following settings were used for viewing the motifs as in our prior publications (Badwal et al., 2020; Higgins et al., 2025): spectrogram view range from 0 Hz to 10,000 Hz, with a dynamic range of 70.0 dB, a pitch setting of 75Hz to 500Hz, and an intensity range of 50 dB to 100 dB. Wav files recorded in SAP were opened consecutively from the start of morning singing until 25 motifs were acquired, to provide sufficient power (Miller et al., 2010). If the same syllable occurred twice within the motif, only the first occurrence of the syllable was selected for analysis. Some syllables were sung frequently within these 25 selected motifs while others were not. This variance was accounted for in our statistical model below. All curated song data files and spectrogram examples for each finch used in this study are found at the University of Arizona ReData repository, available at the time of publication and accessible here: DOI: https://doi.org/10.25422/azu.data.29387402

Syllables were defined as noisy (consonant-like), harmonic (vowel-like), or mixed (harmonic and noisy components) based on their visual appearance on a spectrogram (**Figure 1**) and by CPPS scores, which provide an additional objective assessment, as described in our previous publications (Badwal et al., 2020; Badwal et al., 2018). To summarize, harmonic syllables have higher CPPS scores than noisy syllables and mixed syllables tend to lie in-between. A higher CPPS value means more periodicity in the voice/birdsong. An individual with a hoarse, breathy voice will have a lower CPPS value compared to a non-impaired individual (Brockmann-Bauser et al., 2021; Schultz et al., 2023). Visual inspection reveals that noisy birdsong syllables have lower spectral continuity; their frequency contours are quite variable and their CPPS scores are low. In contrast, harmonic syllables are represented by high spectral continuity indicated by flat stacks with stable periodicity and higher CPPS scores compared to noisy syllables (Kao et al., 2005; Tchernichovski et al., 2000). Mixed syllables comprise both noisy and harmonic notes, have a longer duration (length), with their CPPS scores generally falling in-between noisy and harmonic syllables (Badwal et al., 2020; Badwal et al., 2018). Per aging human voice studies and our prior publications, the following features were extracted from all birdsong syllables using custom code (Supplemental Material: Code Script provided by Dr. Brad Story, University of Arizona): 1) Duration is the length of the syllable from start to finish measured in milliseconds. 2) Mean intensity (or amplitude) in decibels (dB) is perceived as loudness. 3) Cepstral Peak Prominence Smoothed (CPPS) in dB. Smoothing refers to a process that averages several adjacent cepstra before measuring the prominence. The process has the effect of improving how closely CPPS relates to human voice quality (Hillenbrand & Houde, 1996).

3) For harmonic syllables only, measurements were obtained for: mean fundamental frequency (f_o_, pitch in Hertz-Hz), standard deviation of f_o_ (f_o_ std, Hz), and coefficient of variation (CV). CV is calculated in two ways: the CV within a syllable copy (f_o_ std/f_o_ mean) and CV across syllable copies (e.g. renditions) which was calculated by dividing the standard deviation/mean for all copies of that syllable and represents variability over multiple birdsongs (Kao et al., 2005). The f_o_ was not calculated for noisy or mixed birdsong syllables because they lack a consistent f_o_. Our prior publications provide more details concerning acoustic measurements of birdsong and relevance to human voice (Badwal et al., 2020; Badwal et al., 2018). After running the code script in Praat (Supplemental Material), we imported the output values for all measurements into excel spreadsheets which underwent further quality control checks to exclude erroneous values due to technical errors or outliers. Sheets were then imported into the SPSS program (IBM software, version 29) for statistical analyses and plots generated.

### Statistical Analyses

Up to 25 copies per syllable were included using a Generalized Linear Model (GLM) in SPSS. Statistical analyses were conducted in consultation with our University of Arizona statisticians (Acknowledgements). Our prior work indicated no further increase in power beyond n ≥ 25 syllables/motifs in a given behavioral condition sung in a two-hour recording period (Miller et al., 2010). These statistical models are useful for taking into account multiple, non-independent data points when dealing with behavioral data (e.g. syllable types, multiple varied copies, age categories, etc) as in our finch work (Badwal et al., 2020; Higgins et al., 2025). In our prior work, we used a different form of this model, a Linear Mixed Model, to account for a random effect representing different finch subjects in a cross-sectional analyses (Badwal et al., 2020). In this present study, we examined age changes within a finch subject across three individual mornings, so the random effect was not the subject (with the exception of CV scores across motifs, see below). The fixed effects in the model are represented by the interaction of the data with two experimental factors, syllable type (harmonic, mixed, noisy) and age (e.g. 8, 24, 40 months). Our dependent variables were acoustic feature measures: All syllable types were evaluated for changes in duration, intensity, and CPPS. Measurements of f_o_ mean, median, standard deviation (std), and CV are reported for the harmonic syllables only. By not collapsing all data to a single mean score per finch subject when conducting these comparisons, we account for natural variance in syllable values over multiple motifs and consideration by syllable type to detect a meaningful effect of age on song. Residuals generated from raw values inputted into the GLM for duration, intensity, CPPS, and mean f_o_, f_o_ std, and CV within a syllable (e.g. intrasyllable) were examined using histograms to verify symmetrical distribution and validate conformity to the model. Residuals were fairly symmetrical with occasional skew deviating from normal among the three mornings. **Supplemental Table 1** reports the directional change and the post-hoc pairwise comparisons with Bonferroni adjusted p-values for p≤0.05 for all six finches and across the three mornings. Any correlation between syllable type and age was deemed significant if p values ≤0.05. **Supplemental Tables 2-7** represent each individual bird and contain the GLM statistics (Wald Chi-Square value, degrees of freedom, significant p-values) and raw scores for all measures. **Supplemental File ‘Gordon et al Protocol SPSS GLM Stats’** is a how-to protocol for running the statistics.

For the separate evaluation of CV of fundamental frequency (f_o_), known as syllable pitch variability across the 25 motifs, a linear model was performed for each finch subject. Morning was represented as a random intercept, given prior reports of fluctuation in CV within and across multiple mornings (Kao et al., 2005; Wood et al., 2013). Because morning to morning variance was found to be zero, it was excluded. Therefore, a linear model with predictors for age, harmonic syllable type (or just age in the case of only one harmonic syllable being present in that motif), and their interaction were evaluated within individual birds using the open source R language for statistical computing (version 4.3.3). To improve the normality of the residuals, we log10-transformed CV scores for syllables within a finch subject. To compare log-transformed CV scores between birds, we fit a linear mixed effects model with age as a fixed effect and random intercepts for birds and for syllables nested within a bird. We additionally included a random slope for age per syllable nested within bird. To compare log transformed CV scores for each bird in the model of their individual data and across birds in the model with data from multiple birds, we set age as a fixed effect and then random intercepts for finches and for syllables nested within a finch. In the individual birds with multiple syllables, if the interaction of age and syllable was statistically significant, the age comparisons were made for each syllable separately; otherwise, the age comparisons were averaged over syllables. The comparisons across birds used Kenward-Roger degrees of freedom (Kenward and Roger, 1997). A Tukey pairwise comparison of the means between age groups (8 vs. 24 months, 8 vs. 40 months, 24 vs 40 months) was then performed first within individual finch subjects and then across all finches. The alpha-level was set at p<0.05 and adjusted for multiple comparisons (Tukey).

#### Transparency and Openness

The Supplemental Material included with this manuscript contain the individual finch spreadsheets of raw values and statistical output (**Supplemental Tables 1-7**). The code script used to generate the raw song feature values in Praat is also provided as a Supplemental file ‘get_all_cpp_expo.praat‘. The birdsong files extracted and analyzed for this study along with spectrogram examples for each finch are available through the University of Arizona ReData repository at the time of journal publication and are accessible here: DOI: https://doi.org/10.25422/azu.data.29387402

## Results

Out of the seven features examined, intensity and f_o_ mean were the most sensitive to aging across the six finches (**Supplemental Table 1).** Intensity changed across age, with different patterns. For example, **Figure 2** shows that intensity consistently decreased with age across the mornings whereas two other finches showed increased intensity with aging for all three syllable types and mornings (**Figs 3**–**4)**. A fifth bird exhibited the greatest intensity values for the harmonic and mixed syllable types at middle age compared to young and older adulthood (**Fig 5**).

**Fig 2.**
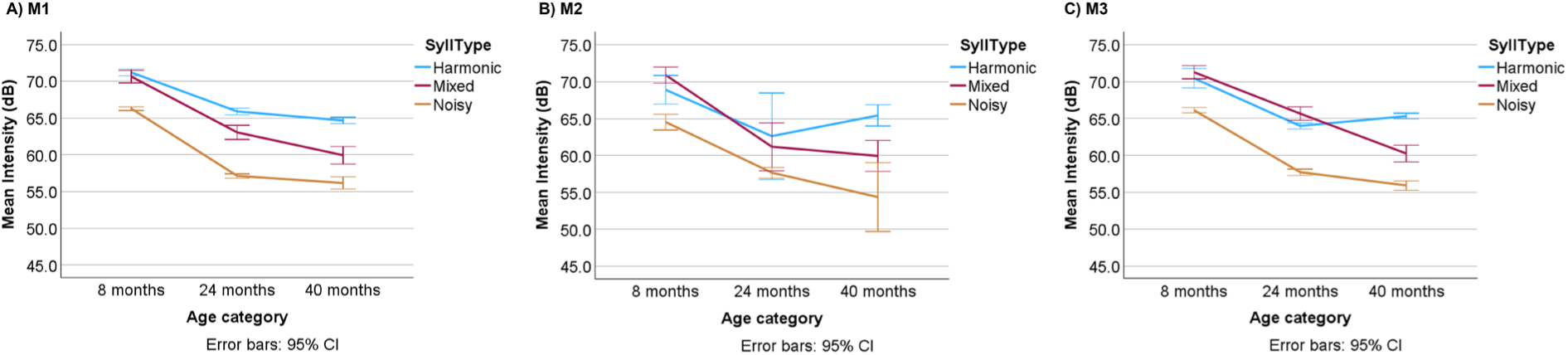
Reduced intensity is pronounced in the transition between young adult and middle age across all three syllable types and at least two mornings. Bird ID: R520 (same as in Figure 1). Plots represent the raw data across the three mornings (M1, M2, M3) with syllable type (SyllType) shown: harmonic syllables (blue), mixed (red), and noisy (dark orange). The age category is represented on the x-axes in months (mon) post-hatch (8-young adult; 24-middle age; 40-older age) per categories defined in the Methods. The song feature (mean intensity) is measured in decibels (dB). Error bars represent the 95% confidence intervals (CI). Pairwise comparisons between the ages were made in our GLM Model with Bonferroni p-value adjustment: **A)** M1: Harmonic and noisy syllables showed significant decreases when comparing 8 v 24 and 8 v 40 mon at p=0.000, while mixed syllables showed decreases across all age comparisons at p=0.000. **B)** M2: No comparisons were significant (p>0.05). **C)** M3: Both harmonic and noisy syllables showed significant decreases when comparing 8 v 24 and 8 v 40 mon, at p=0.000 but not for 24 v 40 mon (p>0.05). Mixed syllables showed significance across all age comparisons (p=0.000).

**Fig 3.**
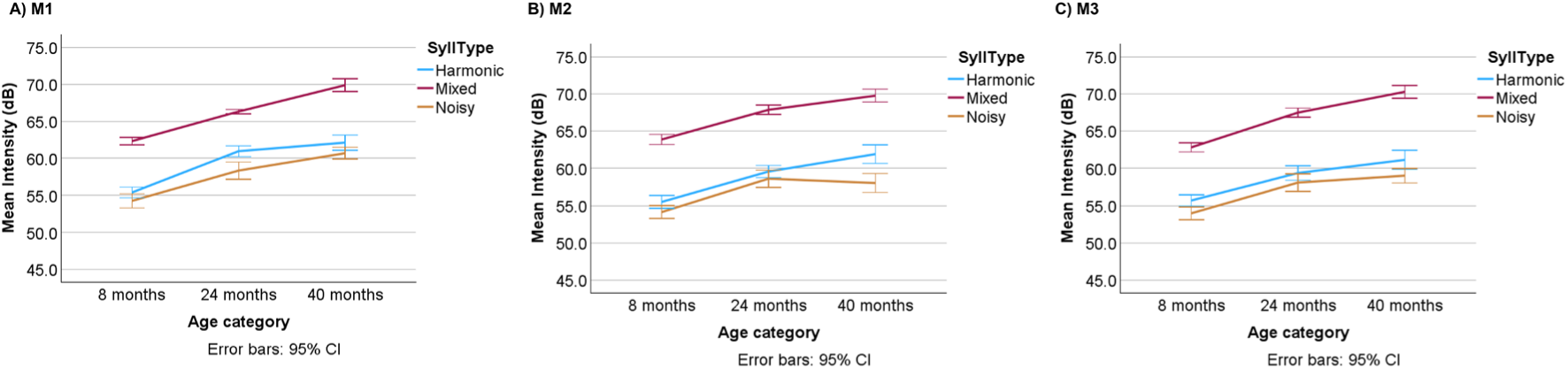
Intensity shows a consistent increase across all three syllable types and mornings as the finch ages. Bird ID: R524. Plots represent the raw data across the three mornings (M1, M2, M3) with syllable type (SyllType) shown: harmonic syllables (blue), mixed (red), and noisy (dark orange). The age category is represented on the x-axes in months (mon) post-hatch (8-young adult; 24-middle age; 40-older age) per categories defined in the Methods. The song feature (mean intensity) is measured in decibels (dB). Error bars represent the 95% confidence intervals (CI). Pairwise comparisons between the ages were made in our GLM Model with Bonferroni p-value adjustment: **A)** M1: Harmonic syllables showed significant increases when comparing 8 v 24 and 8 v 40 mon at p=0.000. Intensity of mixed and noisy syllables increased with aging for all comparisons at p=0.000 with the 24 v 40 mon for mixed syllables at p=0.020. **B)** M2: Harmonic and mixed syllables show increases with aging for all comparisons (harmonic + mixed: 8 v 24 and 8 v 40 mon at p=0.000; Harmonic 24 v 40 mon at p=0.009; Mixed 24 v 40 mon at p=0.013). Noisy syllables increased with aging for 8 v 24 and 8 vs 40 mon (p=0.000) but not for 24 v 40 mon. **C) M3:** Harmonic, mixed, and noisy syllables show increased intensity with aging for 8 v 24 and 8 v 40 mon at p=0.000, but only the mixed syllables show a significant increase at 24 v 40 mon (p=0.000).

**Fig 4.**
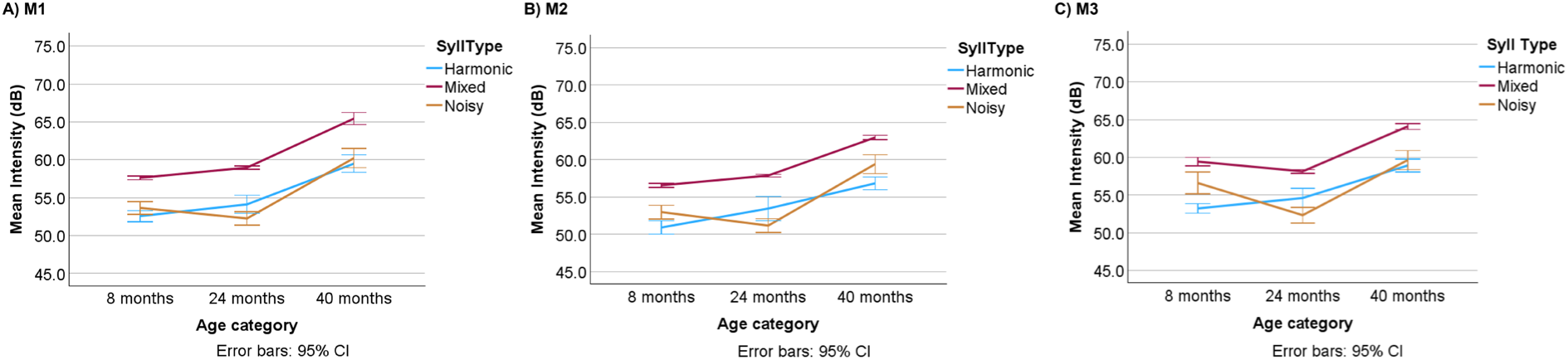
Increased intensity is pronounced in the transition between middle and older ages across all three syllable types and mornings. Bird ID: R526. Plots represent the raw data across the three mornings (M1, M2, M3) with syllable type (SyllType) shown: harmonic syllables (blue), mixed (red), and noisy (dark orange). The age category is represented on the x-axes in months (mon) post-hatch (8-young adult; 24-middle age; 40-older age) per categories defined in the Methods. The song feature (mean intensity) is measured in decibels (dB). Error bars represent the 95% confidence intervals (CI). Pairwise comparisons between the ages were made in our GLM Model with Bonferroni p-value adjustment: **A)** M1: All three syllable types show significant increases in intensity when comparing 24 v 40 and 8 v 40 mon at p=0.000 but no significance for the 8 v 24 mon comparison (p>0.05). **B)** M2: All three syllable types show significant increases in intensity when comparing 24 v 40 and 8 v 40 mon at p=0.000 for mixed and noisy syllables (p=0.001) and for harmonic syllables at 24 v 40 and 8 v 40 mon respectively (p=0.000). The 8 v 24 mon comparison was not significant for any syllable type (p>0.05). **C)** M3: Harmonic syllables showed significant increases for both 24 v 40 and 8 v 40 mon at p=0.000. Mixed syllables showed the same pattern (24 v 40 mon, p=0.000; 8 v 40 mon, p=0.007). Noisy syllables showed reduced intensity when comparing 8 v 24 (p=0.000) followed by an increase from 24 to 40 mon (p=0.000) and for 8 v 40 mon (p=0.001).

**Fig. 5.**
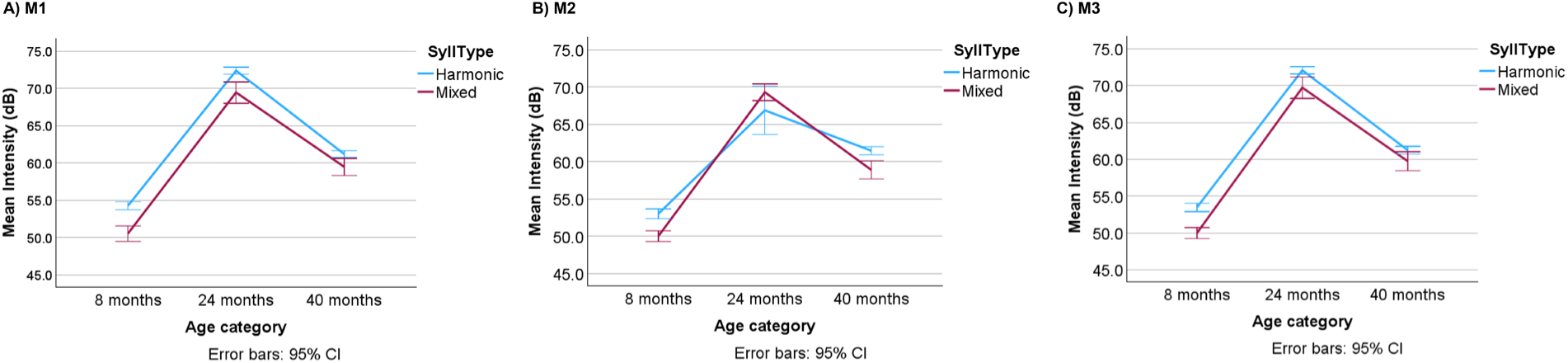
Intensity is highest at middle age across two syllable types and three mornings. Bird ID: R525. Plots represent the raw data across the three mornings (M1, M2, M3) with syllable type (SyllType) shown: harmonic syllables (blue), mixed (red); *this bird lacked noisy syllables in its motif*. The age category is represented on the x-axes in months (mon) post-hatch (8-young adult; 24-middle age; 40-older age) per categories defined in the Methods. The song feature (mean intensity) is measured in decibels (dB). Error bars represent the 95% confidence intervals (CI). Pairwise comparisons between the ages were made in our GLM Model with Bonferroni p-value adjustment: **A-C)** M1-M3: Both harmonic and mixed syllables showed significant increases in intensity from 8 to 24 mon (p=0.000) then intensity dropped from 24 mon to 40 mon (p=0.000). Intensity was also higher at 40 mon compared to 8 mon (p=0.000).

With aging, mean f_o_ increased for harmonic syllables in four out of six finches across the three mornings. The steepest increase occurred from young adult to middle age (**Fig 6**) and included a finch with two harmonic syllables in its motif (**Supplemental Table 1).** For the fifth finch, combining values between the two harmonic syllables in its motif resulted in no significant age comparisons because of differences in f_o_ scores between its syllables A and E (**Fig 7**). When these harmonic syllables were separately evaluated, syllable E showed increased f_o_ mean with aging, while syllable A was inconsistent (**Fig 8**). The sixth finch had three harmonic syllables in its motif - two of the syllables showed decreased f_o_ mean with aging and the third syllable did not. The steepest rise in f_o_ mean occurred from young adult to middle age (**Supplemental Table 1**).

**Fig. 6.**
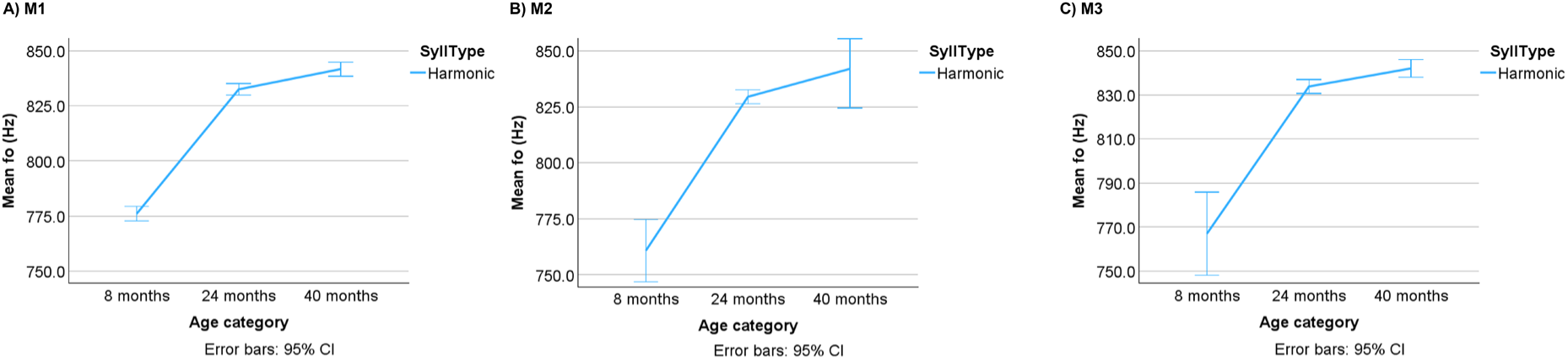
F_o_ mean shows a steep rise from young adult to middle age across all three mornings. Bird ID: R520. Plots represent the raw data across the three mornings (M1, M2, M3) with syllable type (SyllType) showing the only harmonic syllable (blue). The age category is represented on the x-axes in months (mon) post-hatch (8-young adult; 24-middle age; 40-older age) per categories defined in the Methods. The song feature (mean f_o_) is measured in Hertz (Hz). Error bars represent the 95% confidence intervals (CI). Pairwise comparisons between the ages were made in our GLM Model with Bonferroni p-value adjustment: **A)** M1: Comparisons between 8 v 24, 8 v 40, and 24 v 40 mon were all significant (p=0.000). **B-C)** M2, M3: Comparisons between 8 v 24 and 8 v 40 mon were both significant (p=0.000) but 24 v 40 mon was not (p>0.05).

**Fig. 7.**
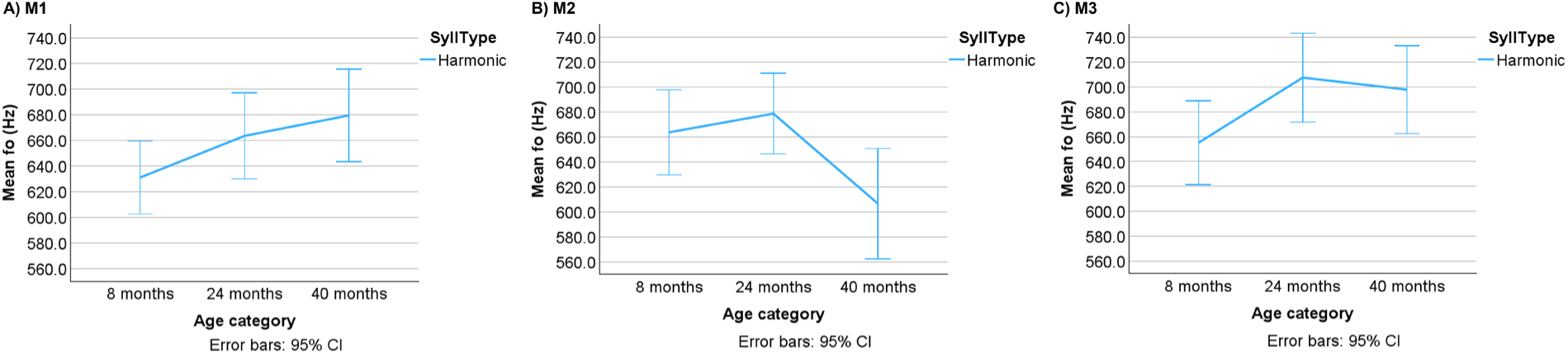
Aging impact on f_o_ mean is muted across the three mornings when scores for two harmonic syllables are combined. Bird ID: R526. Plots represent the raw data across the three mornings (M1, M2, M3) with syllable type (SyllType) showing the average values of syllables A and E together (blue line). The age category is represented on the x-axes in months (mon) post-hatch (8-young adult; 24-middle age; 40-older age) per categories defined in the Methods. The song feature (mean f_o_) is measured in Hertz (Hz). Error bars represent the 95% confidence intervals (CI). Pairwise comparisons between the ages were made in our GLM Model with Bonferroni p-value adjustment: **A-C)** There were no significance differences for the age comparisons in any morning (p>0.05).

**Fig. 8.**
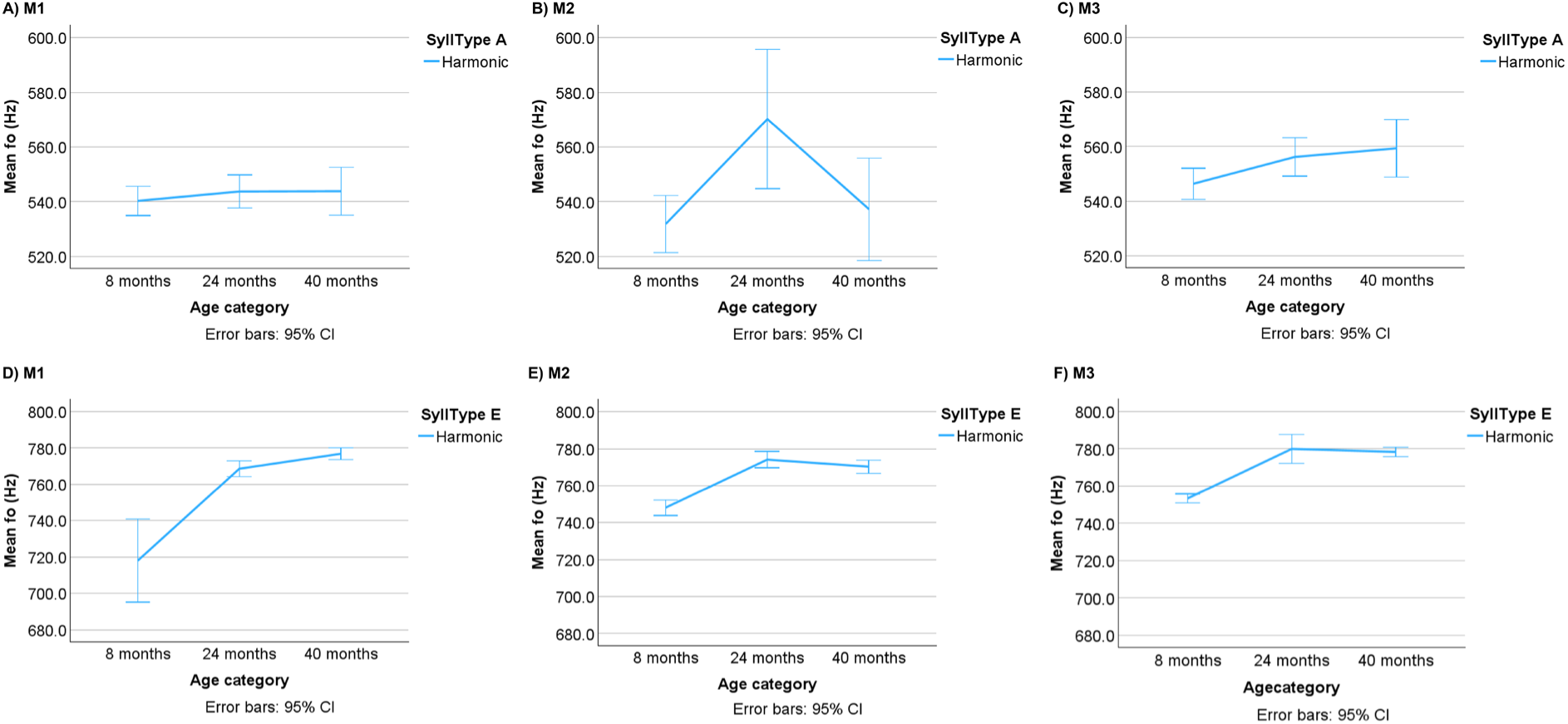
Aging impact on f_o_ mean depends on the individual syllable and morning. Bird ID: R526. Plots represent the raw data across the three mornings (M1, M2, M3) with syllable type (SyllType) showing the individual harmonic syllables A (top row) and E (bottom row). The age category is represented on the x-axes in months (mon) post-hatch (8-young adult; 24-middle age; 40-older age) per categories defined in the Methods. The song feature (mean f_o_) is measured in Hertz (Hz). Error bars represent the 95% confidence intervals (CI). Pairwise comparisons between the ages were made in our GLM Model with Bonferroni p-value adjustment: **A)** M1: Syllable A shows no significant age comparison (p>0.05). **B)** M2: Syllable A increases from 8 to 24 mon (p=0.018) and decreases from 24 to 40 mon (p=0.036) but is not significant for 8 v 40 mon (p>0.05). **C)** M3: Syllable A increases from 8 to 40 mon (p=0.019) but is not significant for the other comparisons (p>0.05). **D-F)** M1, M2, M3: Syllable E increases from 8 to 24 mon and 8 to 40 mon (p=0.000 for both) but not from 24 to 40 mon (p>0.05) across all three mornings.

Two finches showed decreases in intrasyllable f_o_ std and CV scores with aging across all mornings, while the other four finches either exhibited no significant change or inconsistent, syllable-type dependent patterns (**Supplemental Table 1**).

Next, we evaluated CV scores across multiple syllable copies within a bird and across birds (**Supplemental Tables 2-7)**, a standard measurement employed in birdsong research (Kao et al., 2005). No variance in morning to morning variability was found in our model output. There was no statistically significant effect of interaction of age with harmonic syllable (s) within birds or in the model with data from multiple birds (all p > 0.05).

When examining duration and CPPS across all three syllable types, aging effects were much less pronounced. Only two finches showed significant changes in CPPS and duration with age (mainly a decrease for both measures), and these changes were morning and syllable-type dependent (**Supplemental Table 1**).

## Discussion

We employed a longitudinal study design tracking song changes within adult male zebra finches as they aged. Age-related changes in intensity and f_o_ mean represented the most prominent and consistent findings across syllable types and mornings. Our findings also reveal that harmonic syllables should be individually evaluated to take into account their respective patterns when considering an aging impact. Intriguingly, we detected a steep rise in f_o_ mean with age in four finches between young adulthood and middle age with a steep fall in this measure in a fifth finch. Our prior two studies showed no change in f_o_ mean in cross-sectional analyses (Badwal et al., 2020; Higgins et al., 2025), but did show age-related reductions in CPPS and duration. Only two of our finches in this longitudinal study showed changes in these two measures, and they were dependent upon the syllable type and the morning. A potential limitation of our longitudinal study is the sample size - six finches were studied here versus eight to ten finches in our prior cross-sectional analyses (Badwal et al., 2020). However, despite this smaller sample size, the consistency of age-related changes across syllable types and mornings for f_o_ mean suggest that the longitudinal study design is more robust for detecting age-related changes for select measures. It is also important to consider syllable types independently and variability across multiple mornings in any experimental design.

Few longitudinal studies of aging birdsong and human voice exist, and methodologies and experimental design differ across studies. For example, other birdsong studies use different age ranges to define young adult, middle age, and/or old age, sample different numbers of motifs/syllables, may focus only on harmonic syllables, select different acoustic features to evaluate, and/or species (Bengalese instead of Zebra finches) (Cooper et al., 2012; James & Sakata, 2014; Pytte et al., 2007). Despite these differences, age-related song changes were detected. In a study by Pytte et al. (2007), male zebra finch songs were compared over three timepoints within young adulthood (≤15.5 months) and spanning 36-60 months within older adults; middle age was not defined as a separate category. They found an increase in f_o_ mean and decrease in syllable duration within the young adult group timepoints but not in the older adults. In the related Bengalese finch species, Cooper et al. (2012) found that two birds recorded at middle age (spanning one to three years) and then again at old age (six to seven years), showed reduced f_o_ mean, intensity, and greater intersyllable intervals (e.g. a slower song tempo) at old age (Cooper et al., 2012). No change in ventral syringeal muscle fiber composition was detected across multiple birds. James et al. (2014) compared songs in the same Bengalese finches at approximately six months old, which is a comparable age range to our young adult finch cohort, then measured changes again in these same finches at 23 months. The authors reported no change in f_o_ mean, intensity, and duration of harmonic syllables with aging, nor other features. Other syllable types were not examined in their study, but age-related changes in syntax were found (James & Sakata, 2014). Despite species-study differences and age ranges, all studies point to an impact of normative aging on birdsong. This raises the question of how aging targets both central brain mechanisms and the peripheral sound generator, the syrinx. To evaluate these underlying mechanisms, a shared and consistent framework for sampling song, age ranges, and acoustic features needs to be adopted across laboratories.

Data on brain mechanisms involved in aging vocalizations has been largely obtained from studying histological and electrical activity in brainstem-laryngeal circuits in rats (Basken et al., 2012; Bhatt et al., 2025; Johnson et al., 2013; Johnson et al., 2015; Peterson et al., 2013). Work in adult songbird species has mainly focused on on-going neurogenesis in song-dedicated brain nuclei (Aitken et al., 2025; Brenowitz et al., 2024; Hodova et al., 2025; Pytte et al., 2007). As a first step towards uncovering genetic mechanisms underlying aging birdsong, our group discovered that connections between hub genes in finch song nucleus Area X change at middle age and are reduced in old age. Fewer genes are also present in song-associated gene modules (clusters) in the middle and old age categories compared to younger adults (Higgins et al., 2025). How these genetic changes lead to altered neural activity patterns in the song control circuitry in the brain and modulate syringeal control is an open question. Future work in this area would benefit by conducting electrophysiological recordings from Area X and other song nuclei while finches are singing and examine adaptive changes to syringeal muscle physiology (Adam et al., 2021; Adam et al., 2023; Gladman & Elemans, 2024). The ability to quantify song output over the lifetime and directly relate it back to brain or syringeal mechanisms are primary strengths of this songbird model in furthering our understanding of aging voice.

Aging voice studies in humans present multiple challenges due to many confounding factors contributing to variable results across studies. Results vary by task used, such as sustained vowels, reading passages, or conversational speech, as well as by sex (Cavallaro et al., 2024; Rojas et al., 2020; Santos et al., 2023; Schultz et al., 2023; Stathopoulos et al., 2011; Taylor et al., 2020). To compare our findings in aging birdsong to human studies, we focused on f_o_ mean and intensity findings from adult human males only. Consistent with our song data, several studies show a rise in f_o_ mean with age, as reviewed in (Rojas et al., 2020). In a cross-sectional population study, male participants asked to utter a sustained vowel showed a rise in f_o_ mean starting about age 50; intensity also gradually increased with age (Stathopoulos et al., 2011). A metanalysis of a sustained vowel task in a cross-sectional population showed that males 80-89 years had higher f_o_ mean compared to those aged 60-69 years (Rojas et al., 2020). In a longitudinal study of three male speakers providing five minute speech samples, f_o_ mean was found to decrease between 50-70 years of age then rise (Tucker et al., 2023). Intensity is also a robust marker of age-related changes in voice, with older adults showing increases, decreases, or no change in intensity with aging (Dehqan et al., 2013; Huber & SpruilI, 2008; Linville et al., 1989; Rojas et al., 2020; Stathopoulos et al., 2011), a finding also illustrated in our song data. When considering the directional changes (e.g. increases, decreases) in intensity and f_o_ in our aging birdsong dataset, the variability in our findings mirrors those reported in the human subject literature. Despite this variability in the songbird model, the ability to control environmental conditions and subject history along with the advantages of a well-described and experimentally accessible song circuitry highlight the utility of this model for aging research.

## CRediT authorship contribution statement

Michelle L. Gordon: Data Curation, Formal Analysis, Investigation, Methodology, Validation, Visualization, Writing-Original Draft, Writing-review & editing. Arpita Gulati: Data Curation, Formal Analysis, Investigation, Methodology, Validation, Visualization, Writing-review & editing. Robin A. Samlan: Formal Analysis, Validation, Visualization, Writing-original draft, Writing-review & editing. Julie E. Miller: Conceptualization, Data Curation, Formal Analysis, Funding Acquisition, Investigation, Methodology, Project Administration, Resources, Supervision, Validation, Visualization, Writing-original draft, Writing-review & editing.

## ACKNOWLEDGEMENTS

We thank Stephanie J. Munger and the University of Arizona Animal Care staff for finch care. Dr. Brad Story is thanked for his technical input and code. We also thank Dr. Kiana Robinson, Dr. Julia Fisher, and Huashi Li, for their statistical help. Julie E. Miller was supported by startup funds from the Departments of Neuroscience and Speech, Language and Hearing Sciences, University of Arizona, for this project.

## Supplemental Material

### Supplemental Tables

**Table 1. Significant differences for pairwise comparisons with Bonferroni adjustments for acoustic measures across the three age categories and mornings for all six birds.** See embedded legend for further details.

**Table 2. Bird R520.** A) Morning 1, B) Morning 2, C) Morning 3. For each morning, there are five spreadsheets separated as: Tab 1: SPSS Statistical Output from the Generalized Linear Model. Tab 2: Raw Scores from Praat and GLM Residuals from SPSS. Tab 3: Harmonic Raw Scores from Praat and Calculations for CV Within a Syllable. Tab 4: Harmonic Raw Scores from Praat and Calculations for CV Across Syllable Copies

**Table 3. Bird R521.** A) Morning 1, B) Morning 2, C) Morning 3. For each morning, there are five spreadsheets separated as: Tab 1: SPSS Statistical Output from the Generalized Linear Model. Tab 2: Raw Scores from Praat and GLM Residuals from SPSS. Tab 3: Harmonic Raw Scores from Praat and Calculations for CV Within a Syllable. Tab 4: Harmonic Raw Scores from Praat and Calculations for CV Across Syllable Copies.

**Table 4. Bird R535.** A) Morning 1, B) Morning 2, C) Morning 3. For each morning, there are five spreadsheets separated as: Tab 1: SPSS Statistical Output from the Generalized Linear Model. Tab 2: Raw Scores from Praat and GLM Residuals from SPSS. Tab 3: Harmonic Raw Scores from Praat and Calculations for CV Within a Syllable. Tab 4: Harmonic Raw Scores from Praat and Calculations for CV Across Syllable Copies.

**Table 5. Bird R524.** A) Morning 1, B) Morning 2, C) Morning 3. For each morning, there are five spreadsheets separated as: Tab 1: SPSS Statistical Output from the Generalized Linear Model. Tab 2: Raw Scores from Praat and GLM Residuals from SPSS. Tab 3: Harmonic Raw Scores from Praat and Calculations for CV Within a Syllable. Tab 4: Harmonic Raw Scores from Praat and Calculations for CV Across Syllable Copies.

**Table 6. R526.** A) Morning 1, B) Morning 2, C) Morning 3. For each morning, there are five spreadsheets separated as: Tab 1: SPSS Statistical Output from the Generalized Linear Model. Tab 2: Raw Scores from Praat and GLM Residuals from SPSS. Tab 3: Harmonic Raw Scores from Praat and Calculations for CV Within a Syllable. Tab 4: Harmonic Raw Scores from Praat and Calculations for CV Across Syllable Copies.

**Table 7. R525.** A) Morning 1, B) Morning 2, C) Morning 3. For each morning, there are five spreadsheets separated as: Tab 1: SPSS Statistical Output from the Generalized Linear Model. Tab 2: Raw Scores from Praat and GLM Residuals from SPSS. Tab 3: Harmonic Raw Scores from Praat and Calculations for CV Within a Syllable. Tab 4: Harmonic Raw Scores from Praat and Calculations for CV Across Syllable Copies.

**Praat Code Script ‘get_all_cpp_expo.praat.** Script provided by Dr. Brad Story, University of Arizona, for use in Praat to extract features presented in this manuscript and additional features.

**‘Gordon et al Protocol SPSS GLM Stats’.** Powerpoint file provided by Julie E. Miller as a ‘how-to’ guide for running statistics and plots on aging birdsong data in SPSS v29.

**Figure 1 audio files.** Three .wav files representing an example motif from Bird ID R520 at three ages (8 months, 24, 40 months). The file can be played in Windows Media Player, and/or played and visualized in Praat (https://www.fon.hum.uva.nl/praat/) or Audacity software (https://www.audacityteam.org/).

