## Supplementary material for "Voice Changes in Aging Birdsong: A Longitudinal Study in Adult Male Zebra Finches": get_all_cpp_expo.praat

```
# 
# Praat Script
# Get int, dur, foMean, foStd, foMin, foMax, hnr, cpps of all selected sounds
#
# B. Story
# June 2021
#
#
#
clearinfo

#the pause could be commented out if annoying
pause select all sounds to be used for this operation

numberOfSelectedSounds = numberOfSelected ("Sound")

for thisSelectedSound to numberOfSelectedSounds
	sound'thisSelectedSound' = selected("Sound",thisSelectedSound)
endfor

print File       Int.(dB)  Dur.(s) foMedian(Hz) foMean(Hz) foStd(Hz) foMin(Hz) foMax(Hz) foRange(Hz) hnr(dB)  cpp(dB)  cpps(dB) 'newline$' 'newline$'

for thisSound from 1 to numberOfSelectedSounds
    select sound'thisSound'
	name$ = selected$("Sound")


		# extract intensity 
    	##################################################################
    	int = Get intensity (dB)

		# extract duration 
    	##################################################################
		dur = Get total duration

		# extract F0  
    	##################################################################
    	select sound'thisSound'
		name$ = selected$("Sound")
		minPitch = 75
		ceilPitch = 1600
    	timeStep = 0.0
    	#select Sound 'fileName$'
    	To Pitch (cc)... timeStep minPitch 15 1 0.03 0.45 0.01 0.35 0.14 ceilPitch
    	foMean = Get mean... 0 0 Hertz
		foMedian = Get quantile... 0 0 0.50 Hertz
		foStd = Get standard deviation... 0 0 Hertz
		foMin = Get minimum... 0 0 Hertz parabolic
		foMax = Get maximum... 0 0 Hertz parabolic
		foRange = foMax - foMin
		

		# extract harmonic to noise ratio 
    	##################################################################
    	select sound'thisSound'
		name$ = selected$("Sound")
		minPitch = 75
    	timeStep = 0.01
    	To Harmonicity (cc)... timeStep minPitch 0.1 1.0
    	hnr = Get mean... 0 0


		#extract cpp
    	##################################################################
    	select sound'thisSound'
		name$ = selected$("Sound")
		#minPitch determines the window length (3 pitch periods/minPitch)
		minPitch = 300
    	timeStep = 0.002
		maxFreq = 20000
    	To PowerCepstrogram... minPitch timeStep maxFreq 50
    	cpp = Get CPPS... 1 0.0 0.0 300 1500 0.05 parabolic 0.0005 0 "Exponential decay" Robust
		

		#extract cpps 
    	##################################################################
    	select sound'thisSound'
		name$ = selected$("Sound")
		#minPitch determines the window length (3 pitch periods/minPitch)
		minPitch = 300
    	timeStep = 0.002
		maxFreq = 20000
    	To PowerCepstrogram... minPitch timeStep maxFreq 50
		cpps = Get CPPS... 1 0.0 0.0001 300 1500 0.05 parabolic 0.0005 0 "Exponential decay" Robust

		

		# write data to info window
    	print 'name$' 'int:2'   'dur:3'     'foMedian:1'        'foMean:1'       'foStd:1'       'foMin:1'     'foMax:1'   'foRange:1'         'hnr:2'     'cpp:2'   'cpps:2' 'newline$'


   
endfor

#re-select the sounds
select sound1
for thisSound from 2 to numberOfSelectedSounds
    plus sound'thisSound'
endfor
```
