## Supplementary material for "Voice Changes in Aging Birdsong: A Longitudinal Study in Adult Male Zebra Finches": Gordon et al Protocol SPSS GLM Stats.pptx

### Slide 1
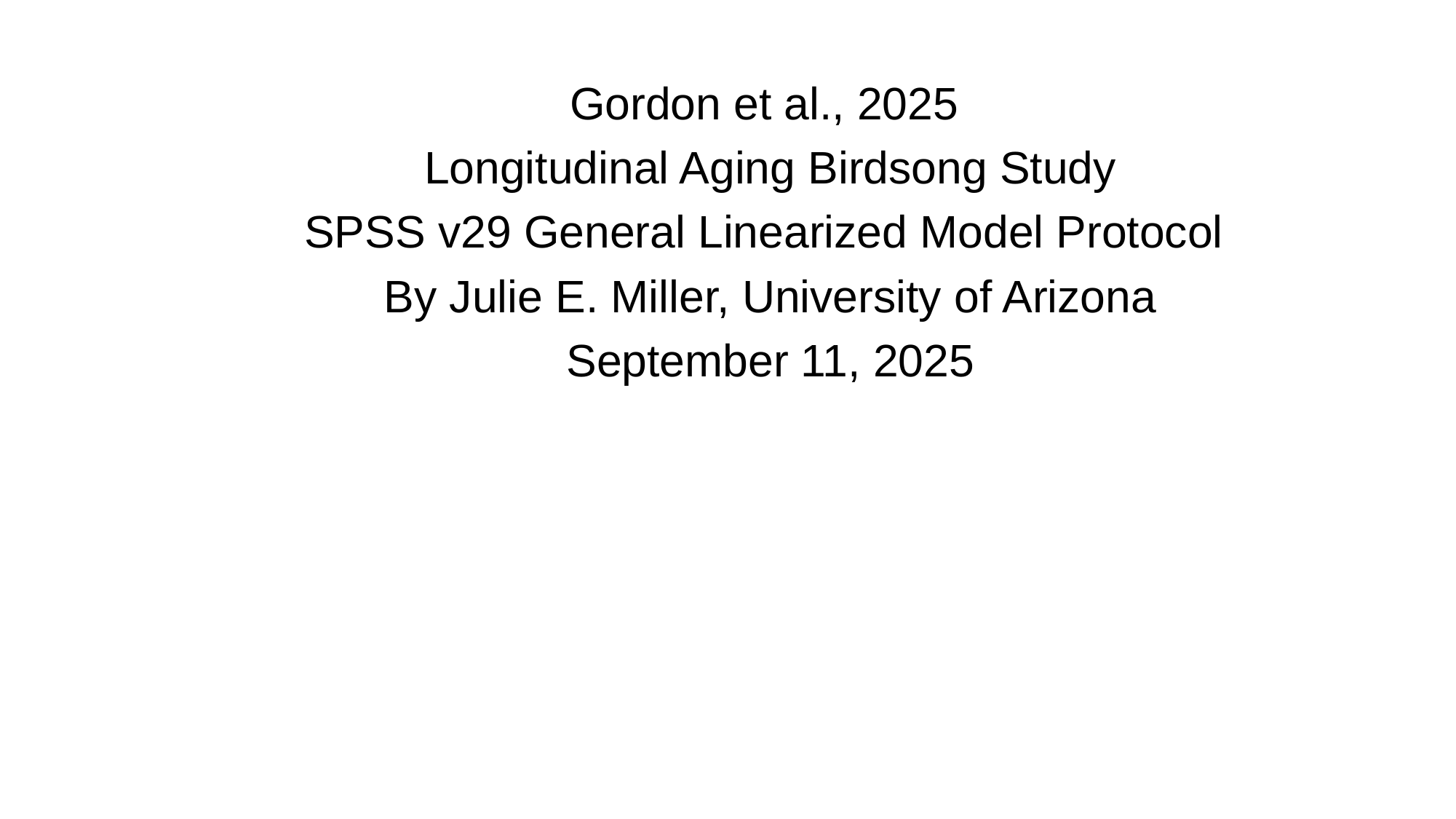

Gordon et al., 2025
Longitudinal Aging Birdsong Study
SPSS v29 General Linearized Model Protocol
By Julie E. Miller, University of Arizona
September 11, 2025

### Slide 2
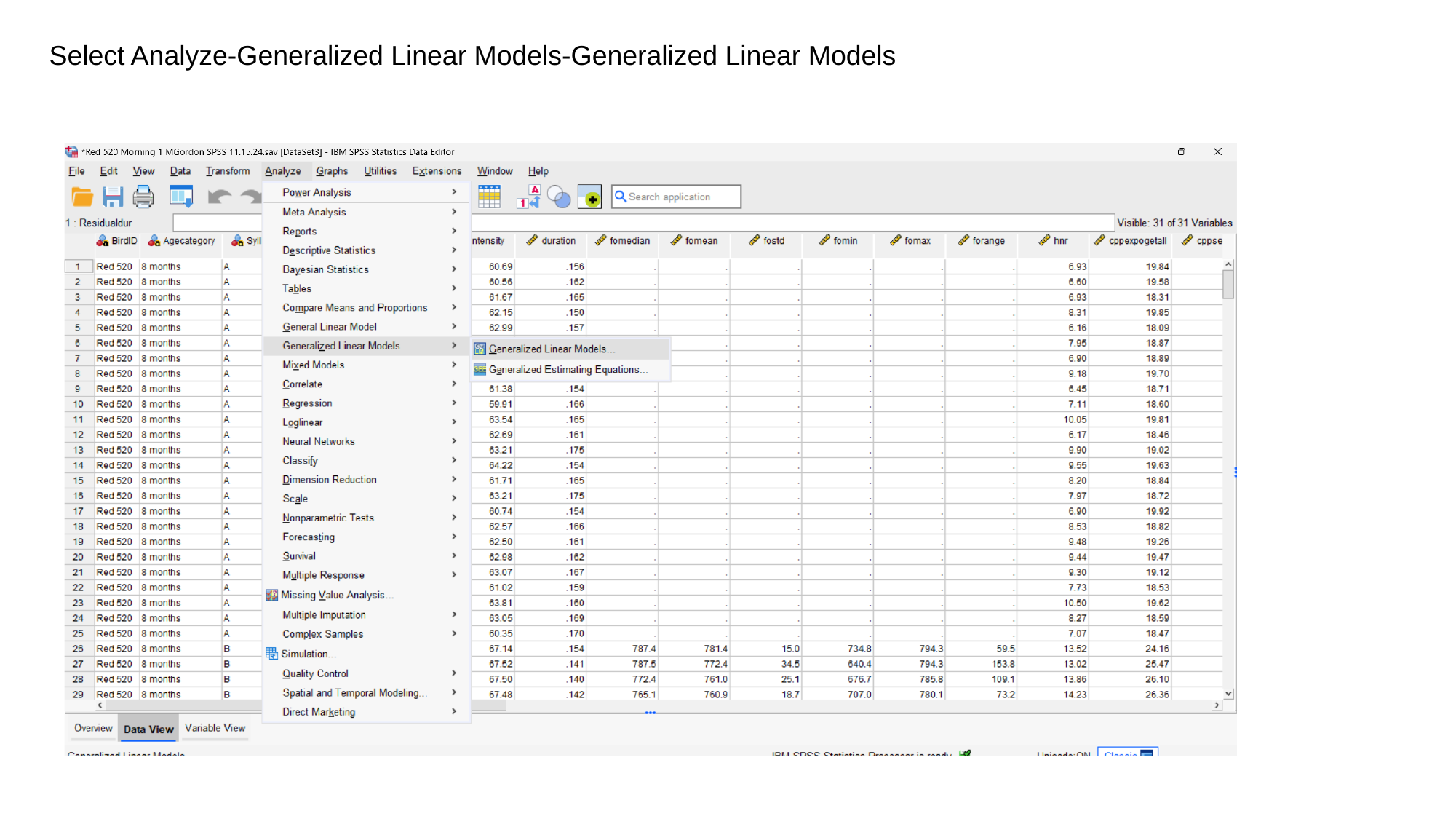

Select Analyze-Generalized Linear Models-Generalized Linear Models

### Slide 3
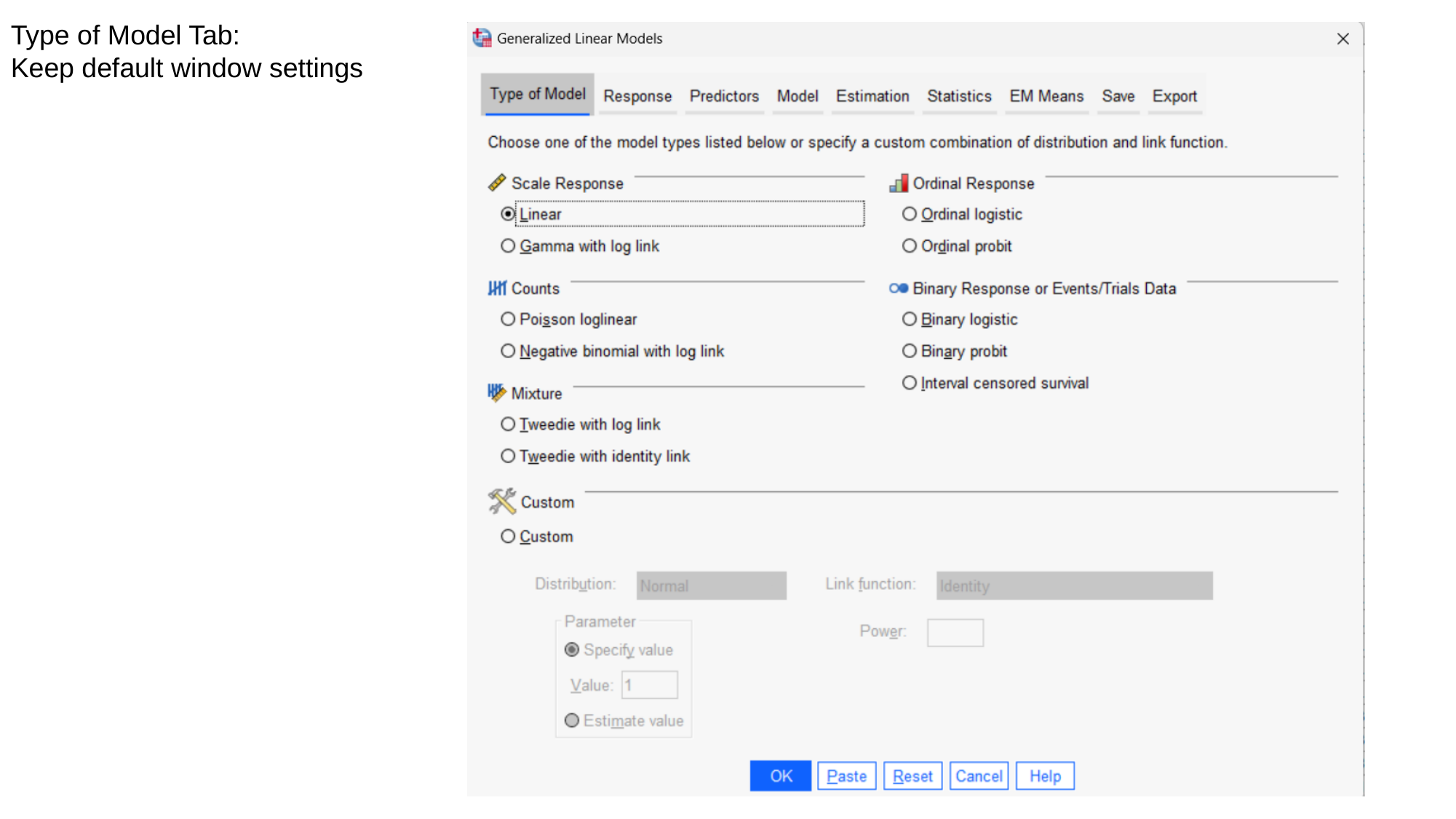

Type of Model Tab:
Keep default window settings

### Slide 4
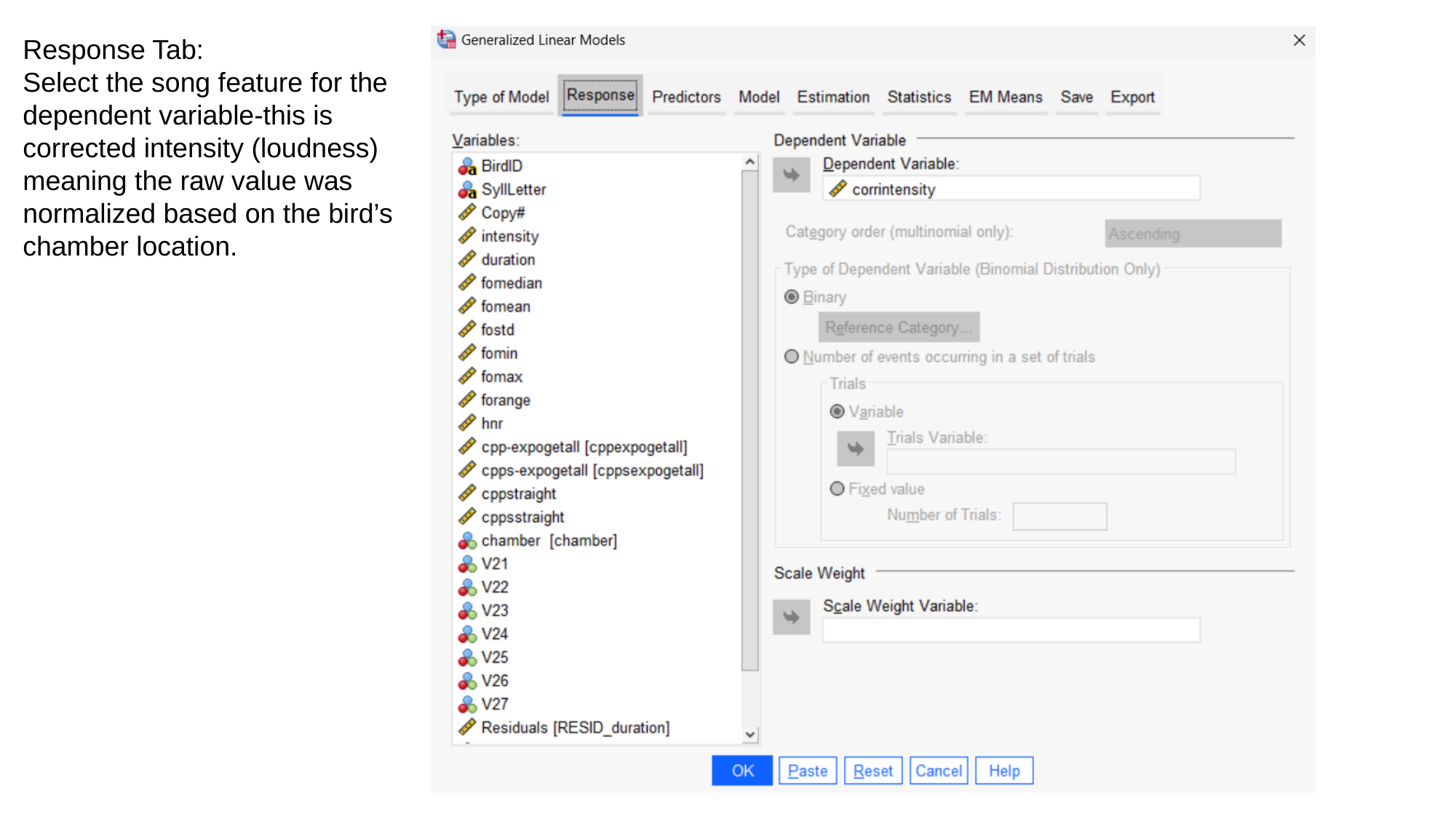

Response Tab:
Select the song feature for the
dependent variable-this is
corrected intensity (loudness) meaning the raw value was normalized based on the bird’s chamber location.

### Slide 5
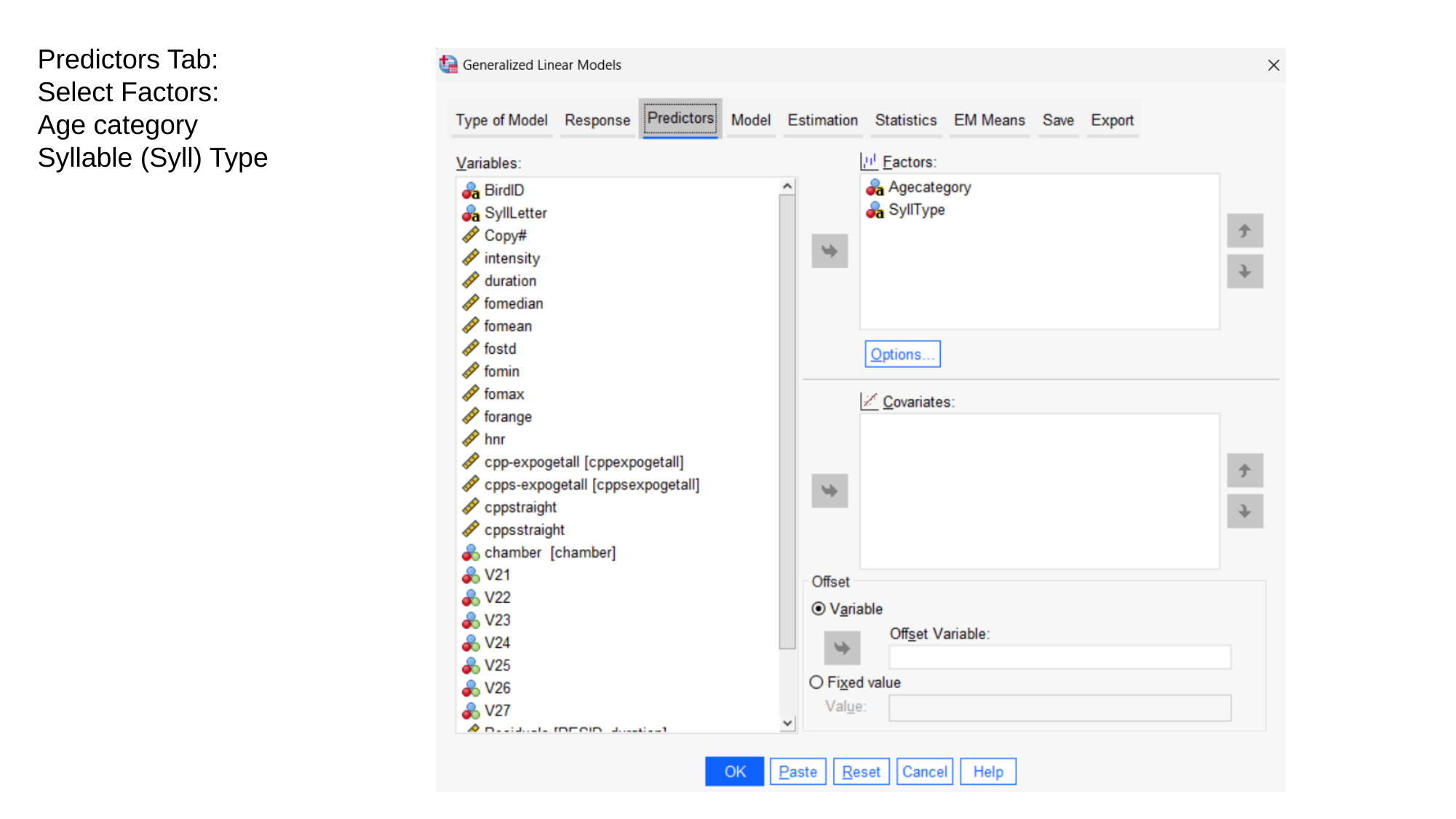

Predictors Tab:
Select Factors:
Age category
Syllable (Syll) Type

### Slide 6
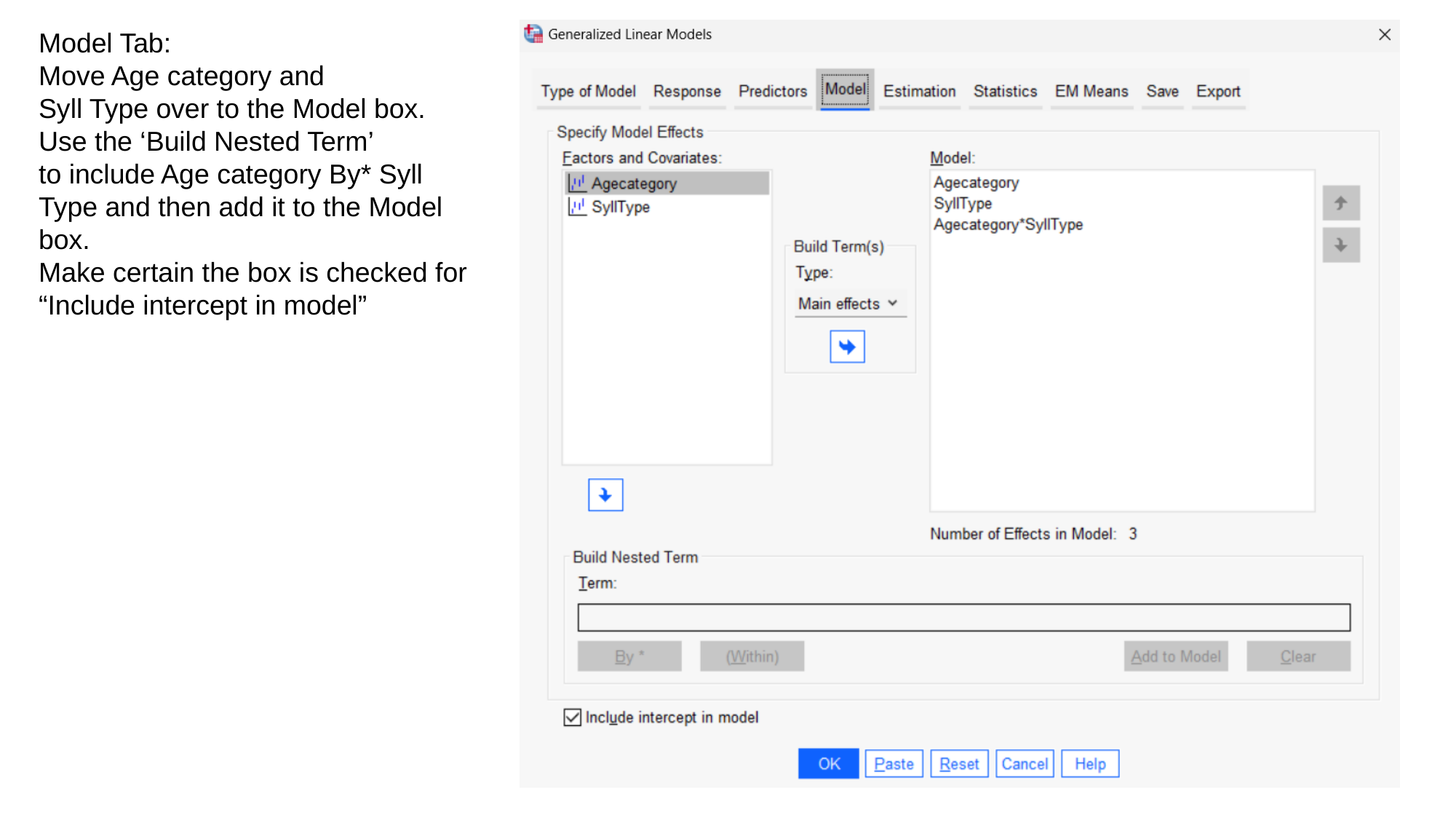

Model Tab:
Move Age category and
Syll Type over to the Model box.
Use the ‘Build Nested Term’
to include Age category By* Syll
Type and then add it to the Model
box.
Make certain the box is checked for
“Include intercept in model”

### Slide 7
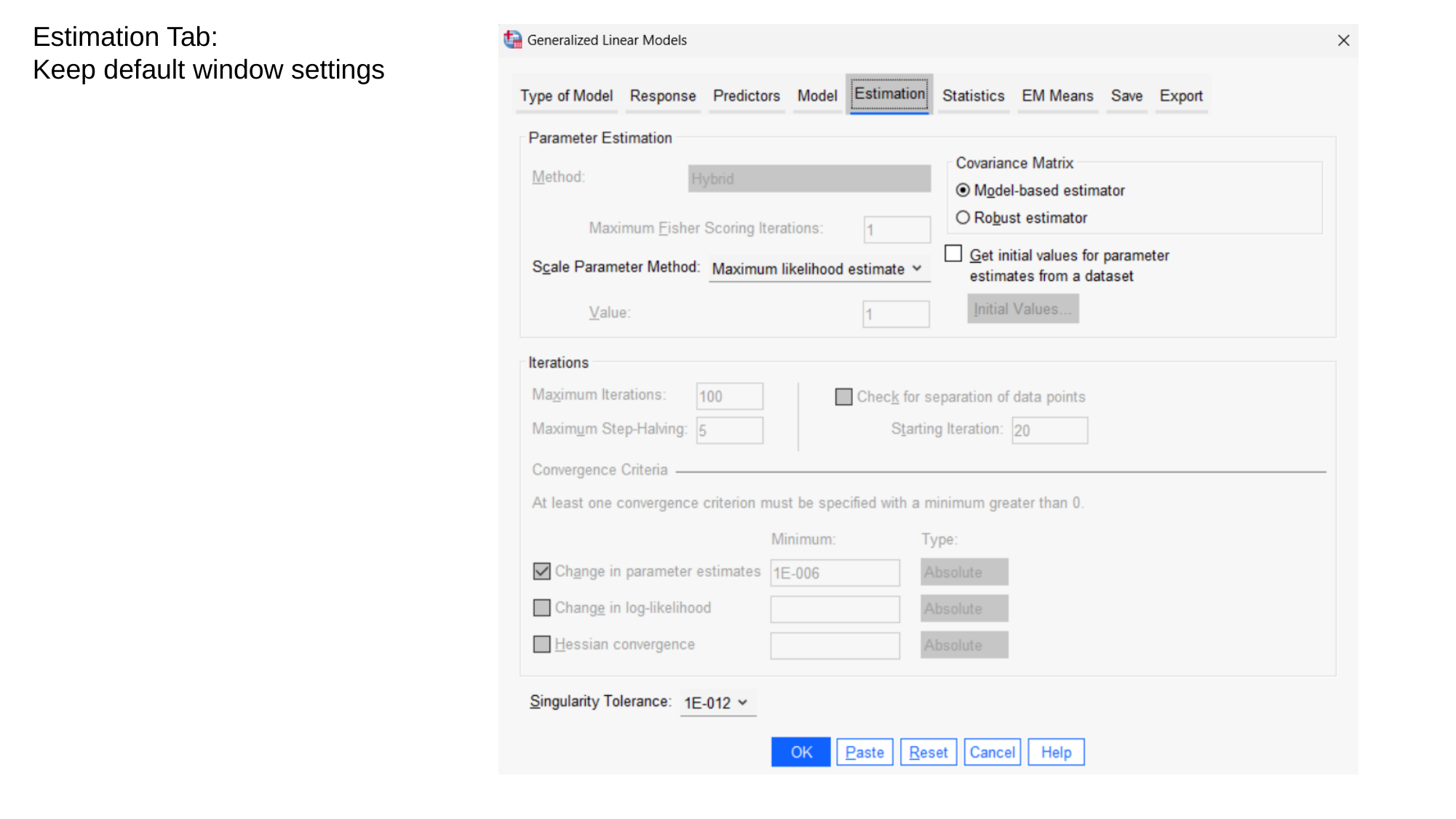

Estimation Tab:
Keep default window settings

### Slide 8
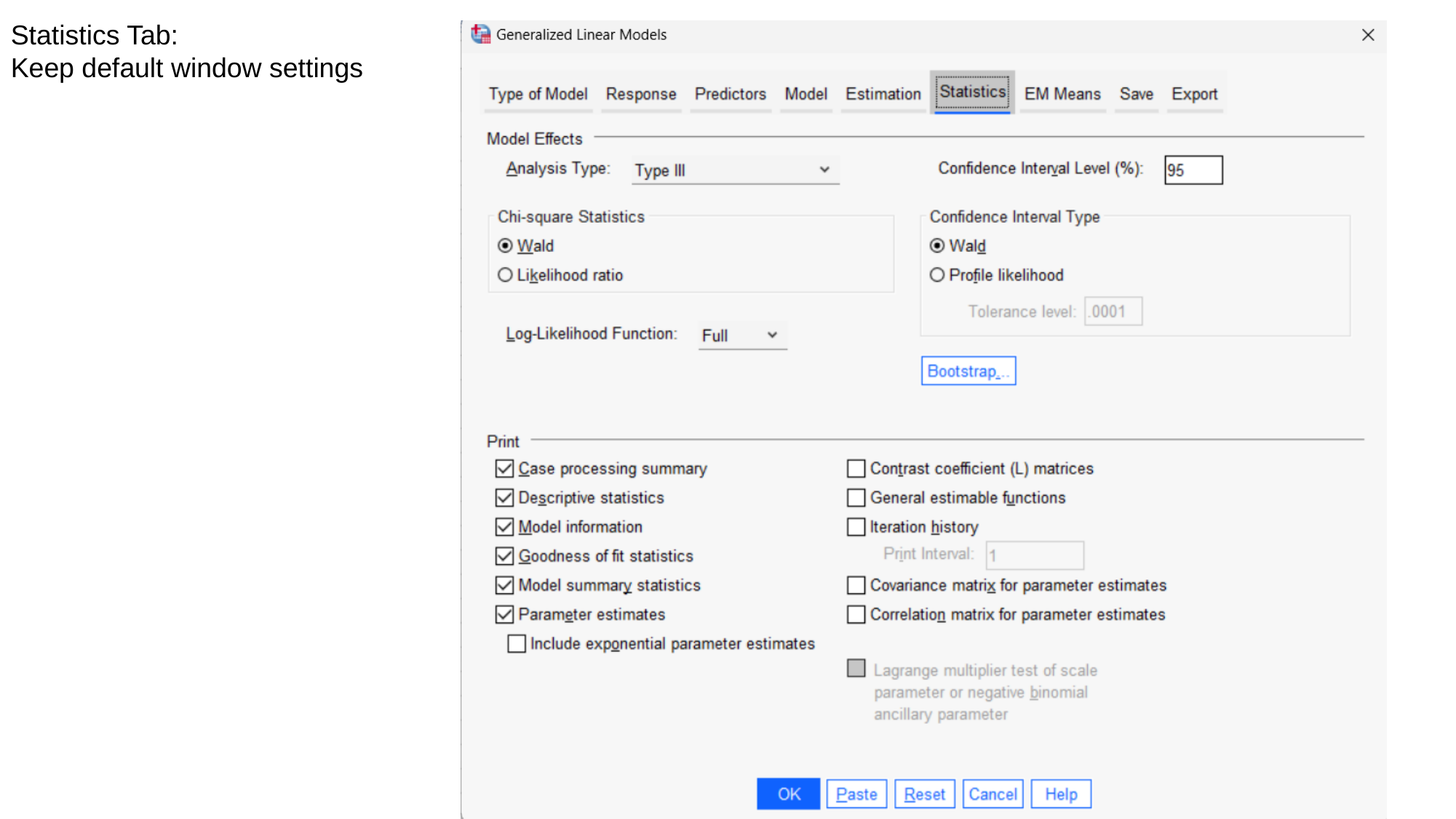

Statistics Tab:
Keep default window settings

### Slide 9
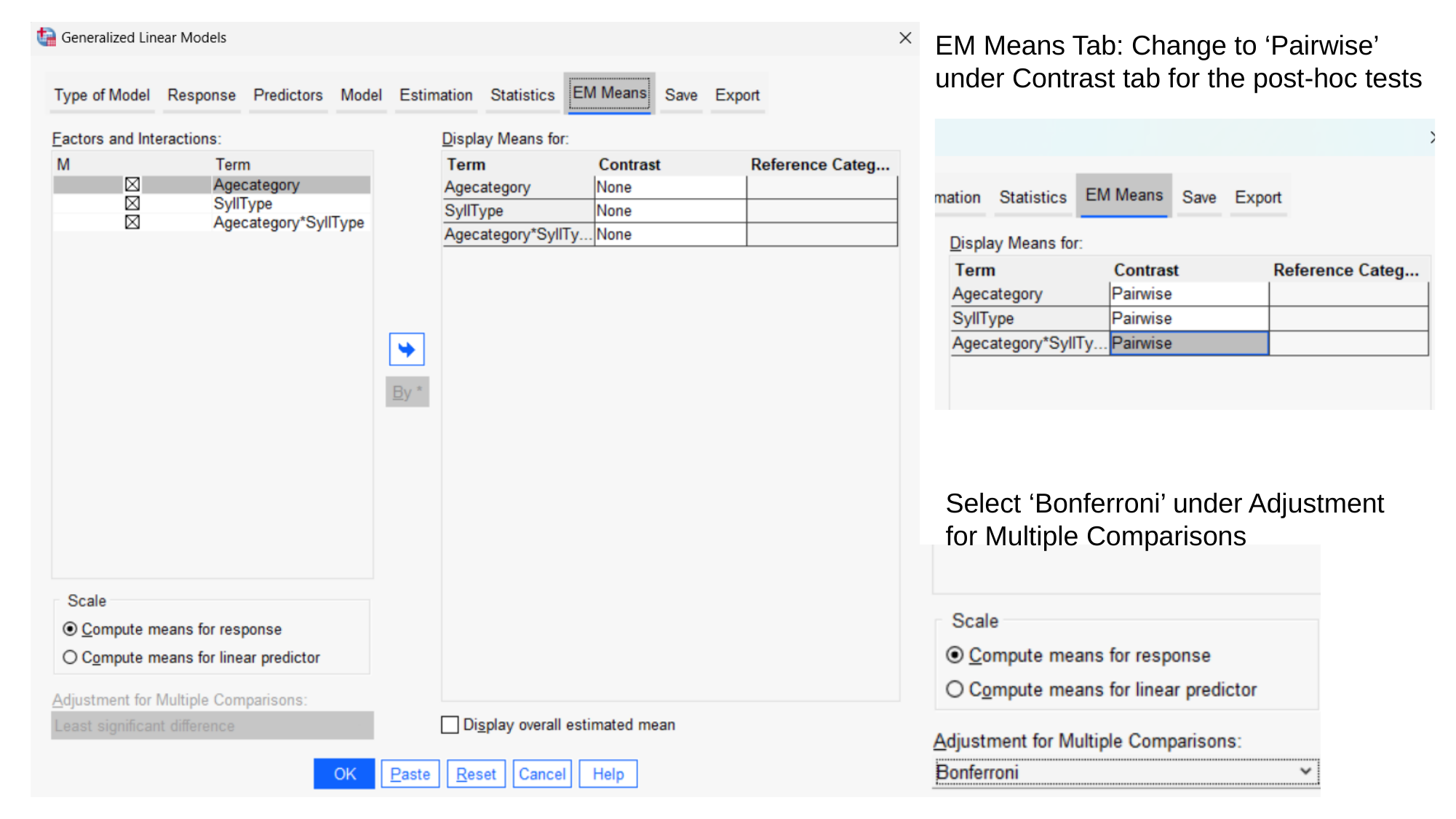

EM Means Tab: Change to ‘Pairwise’ under Contrast tab for the post-hoc tests
Select ‘Bonferroni’ under Adjustment
for Multiple Comparisons

### Slide 10
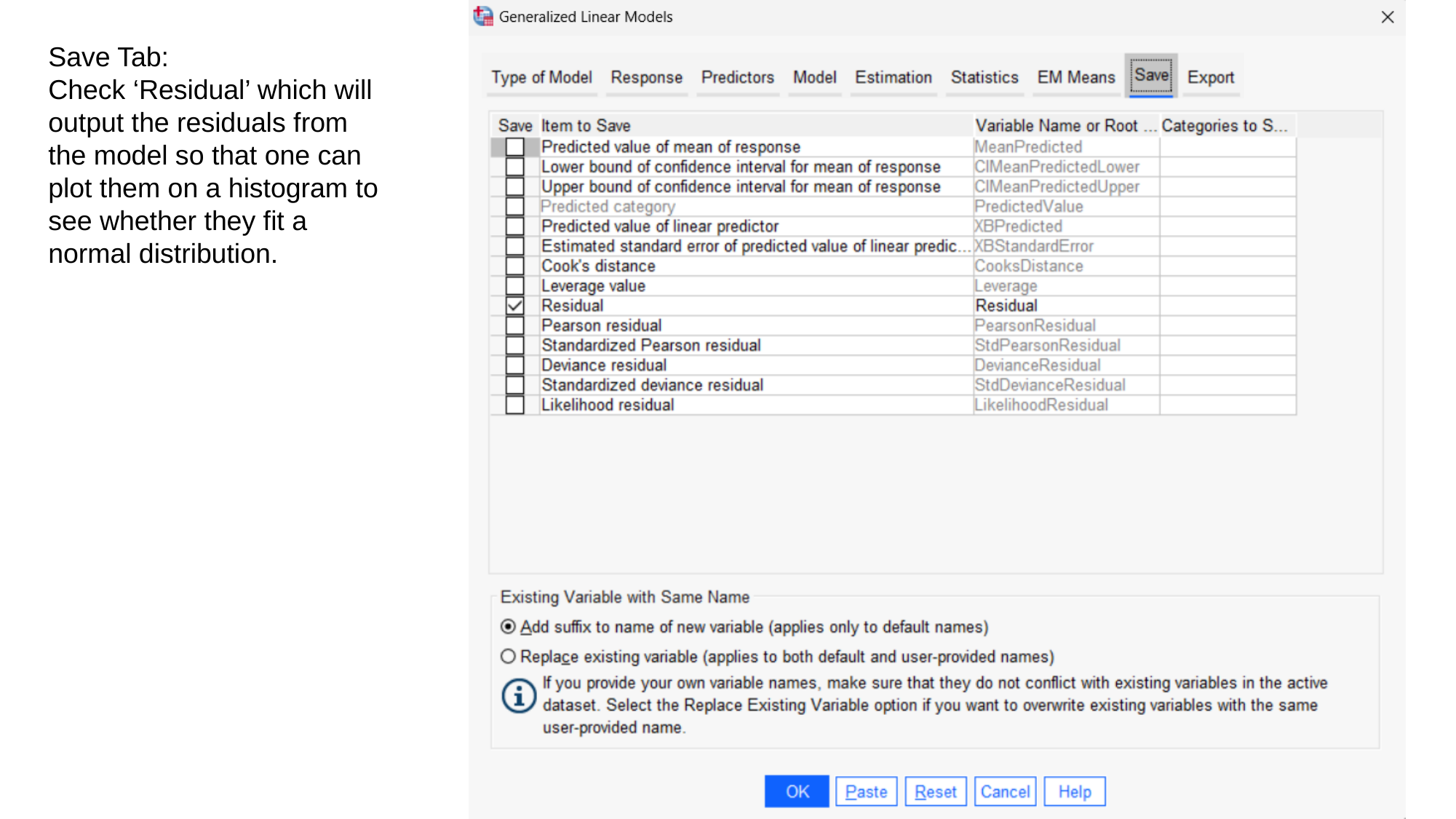

Save Tab:
Check ‘Residual’ which will output the residuals from the model so that one can plot them on a histogram to see whether they fit a normal distribution.

### Slide 11
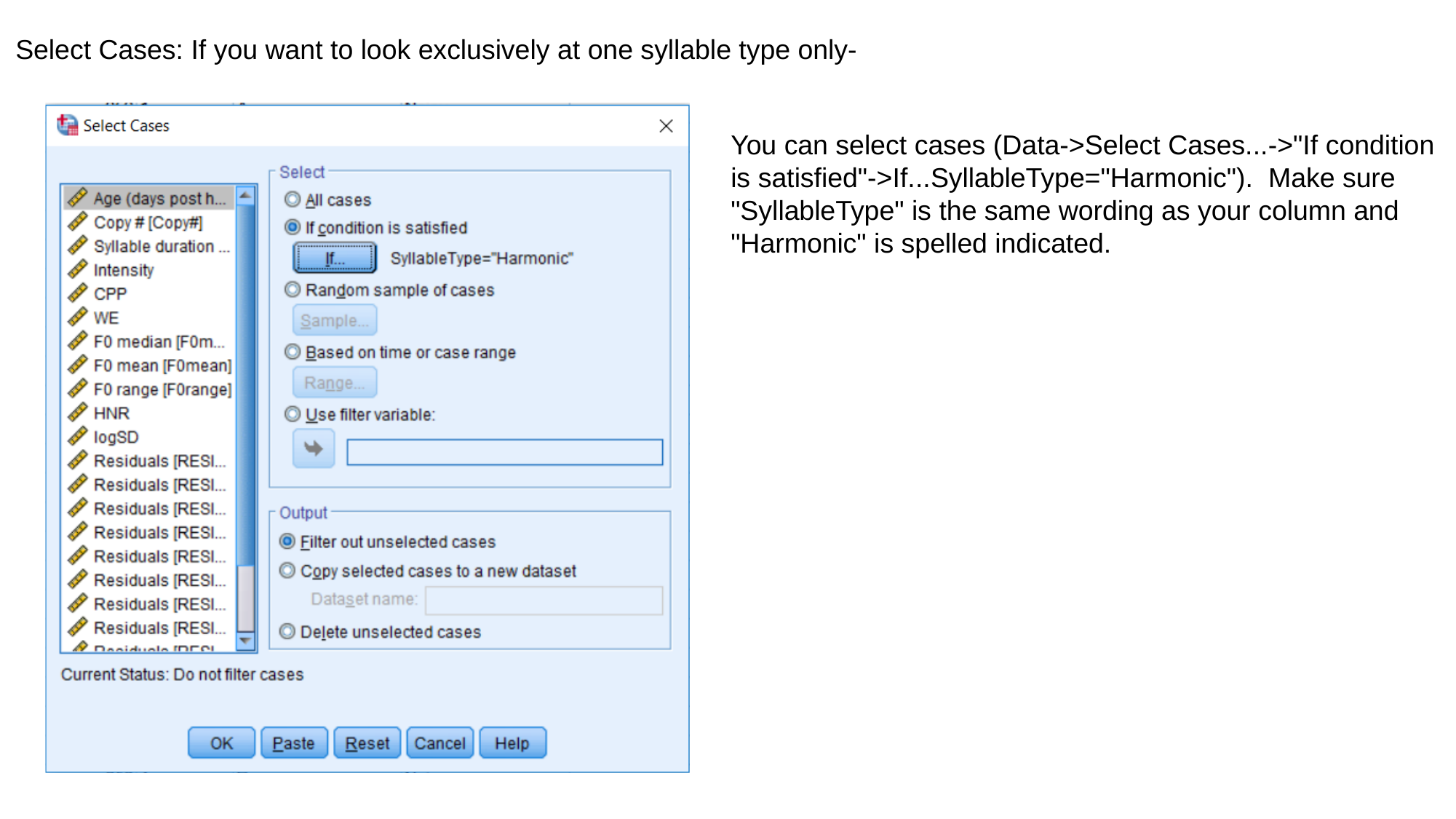

Select Cases: If you want to look exclusively at one syllable type only-
You can select cases (Data->Select Cases...->"If condition is satisfied"->If...SyllableType="Harmonic").  Make sure "SyllableType" is the same wording as your column and "Harmonic" is spelled indicated.

### Slide 12
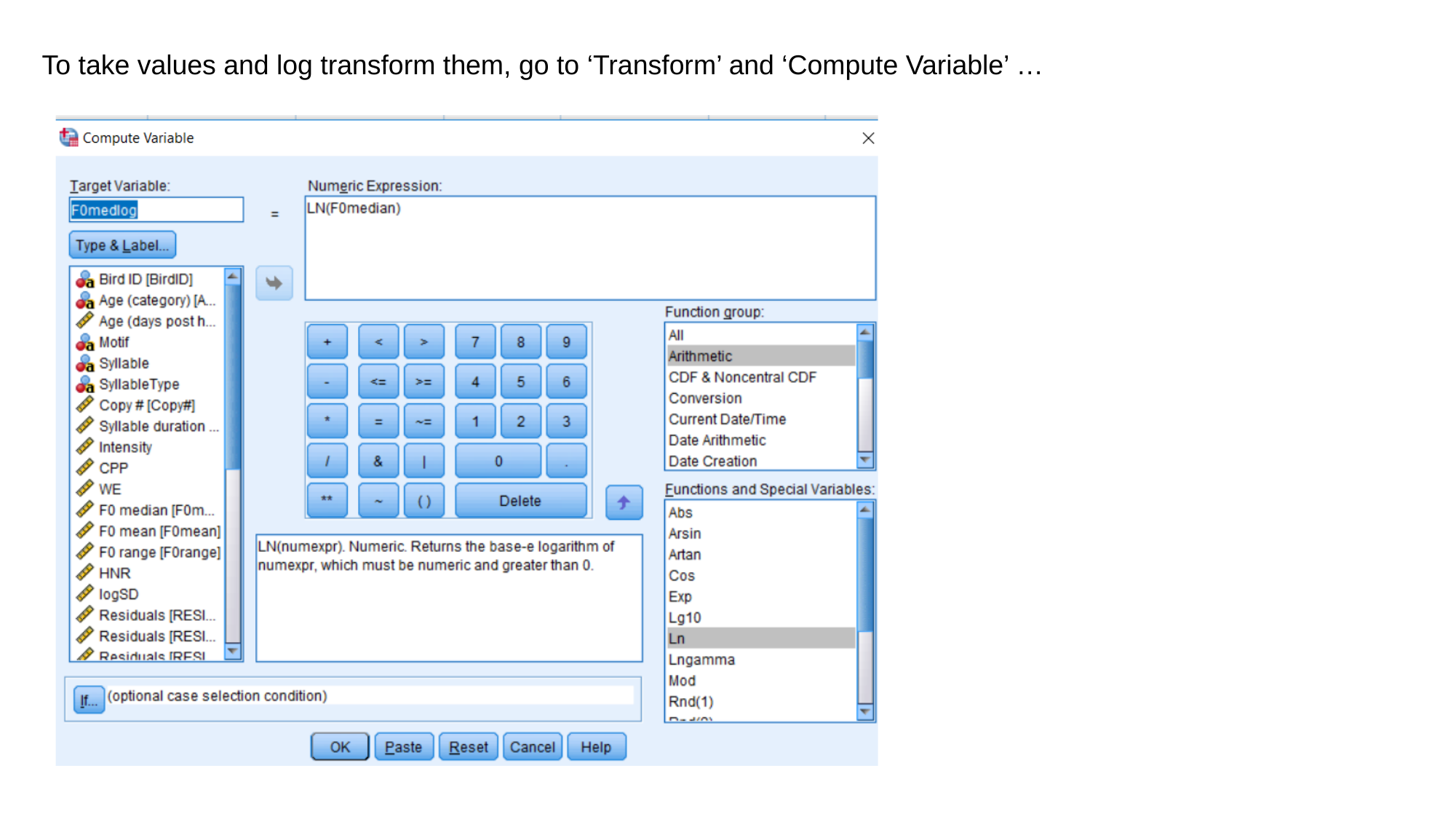

To take values and log transform them, go to ‘Transform’ and ‘Compute Variable’ …

### Slide 13
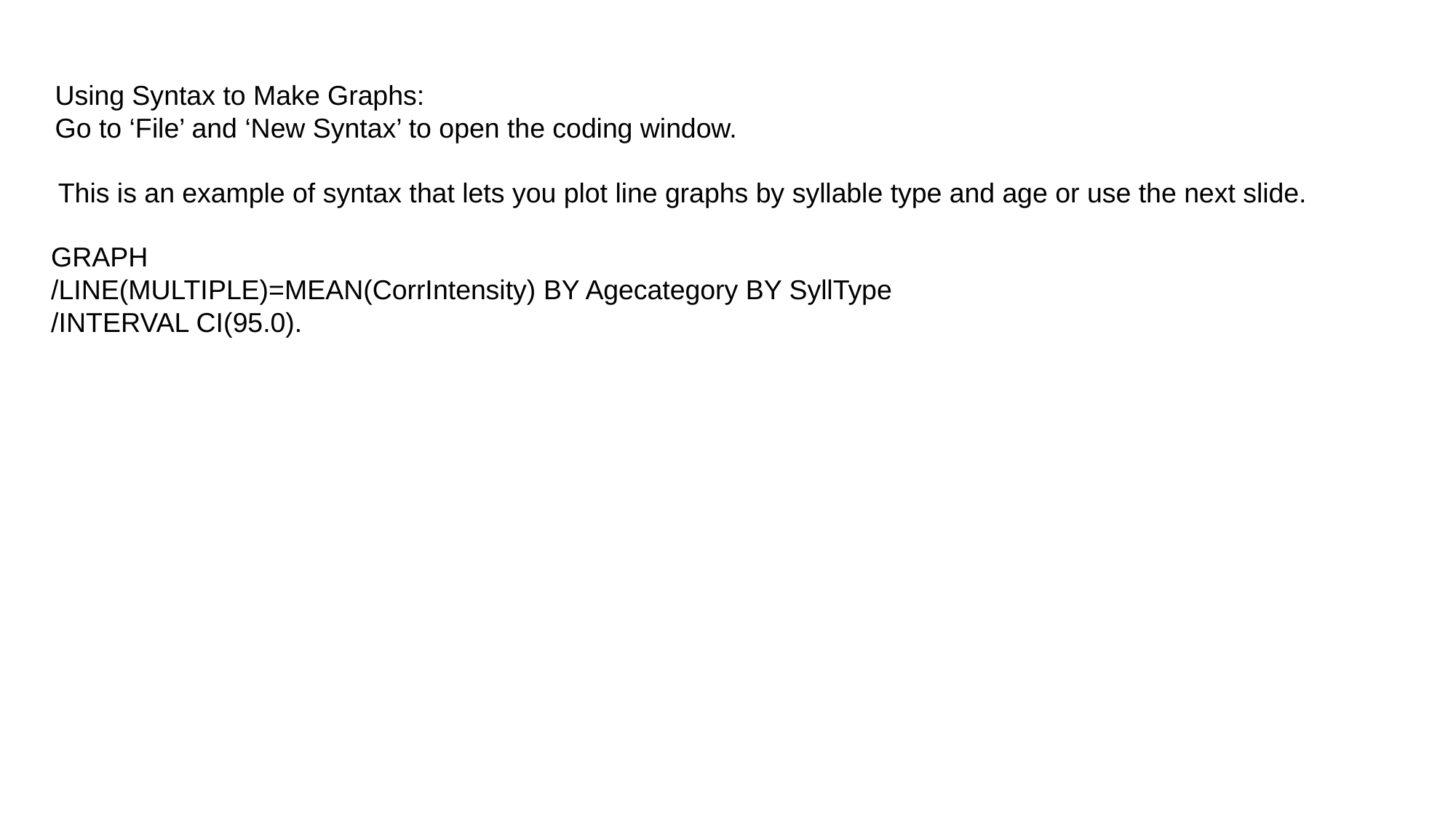

Using Syntax to Make Graphs:
Go to ‘File’ and ‘New Syntax’ to open the coding window.
This is an example of syntax that lets you plot line graphs by syllable type and age or use the next slide.
GRAPH
/LINE(MULTIPLE)=MEAN(CorrIntensity) BY Agecategory BY SyllType
/INTERVAL CI(95.0).

### Slide 14
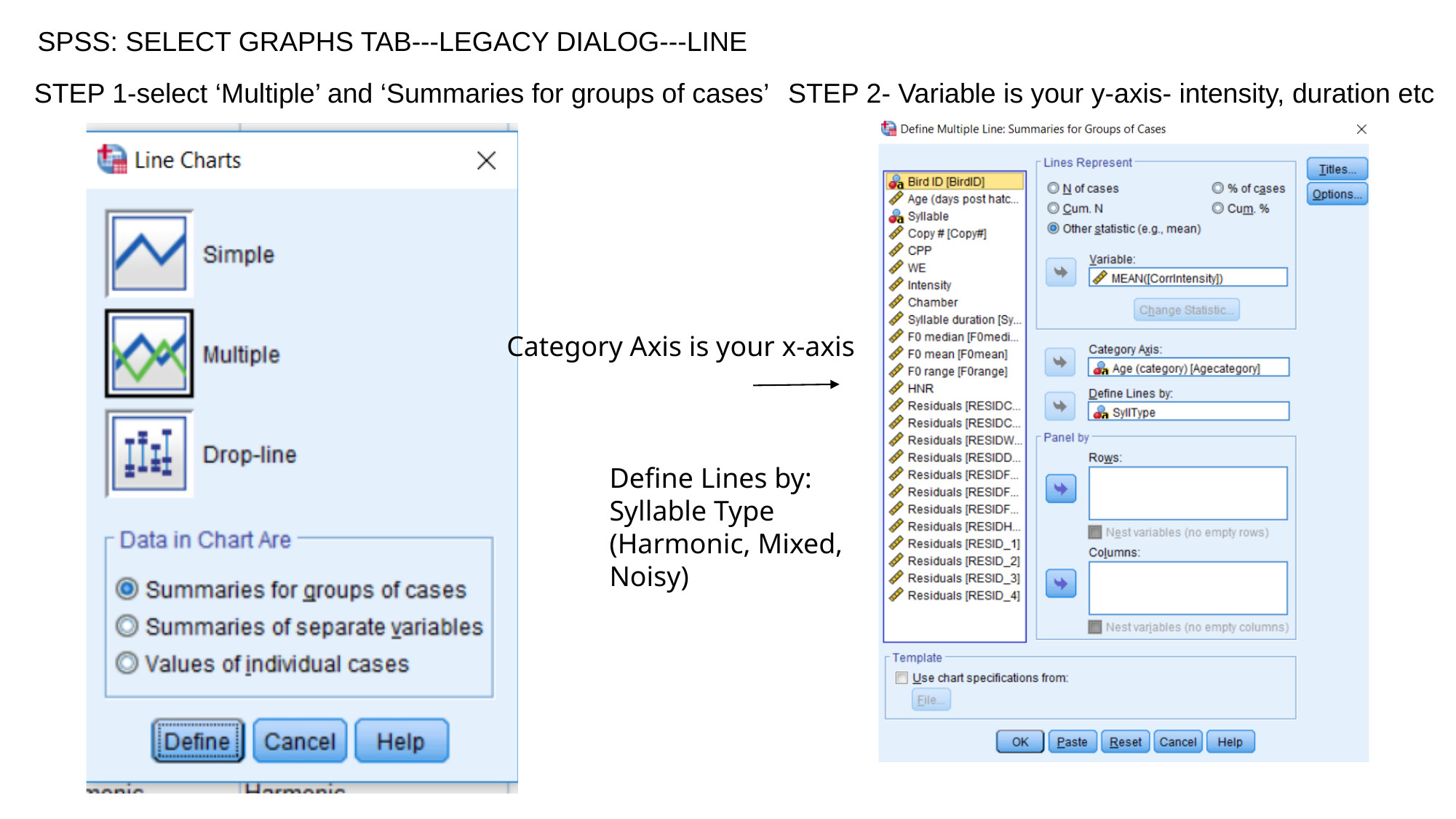

SPSS: SELECT GRAPHS TAB---LEGACY DIALOG---LINE
STEP 1-select ‘Multiple’ and ‘Summaries for groups of cases’
STEP 2- Variable is your y-axis- intensity, duration etc
Category Axis is your x-axis
Define Lines by:
Syllable Type
(Harmonic, Mixed,
Noisy)

### Slide 15
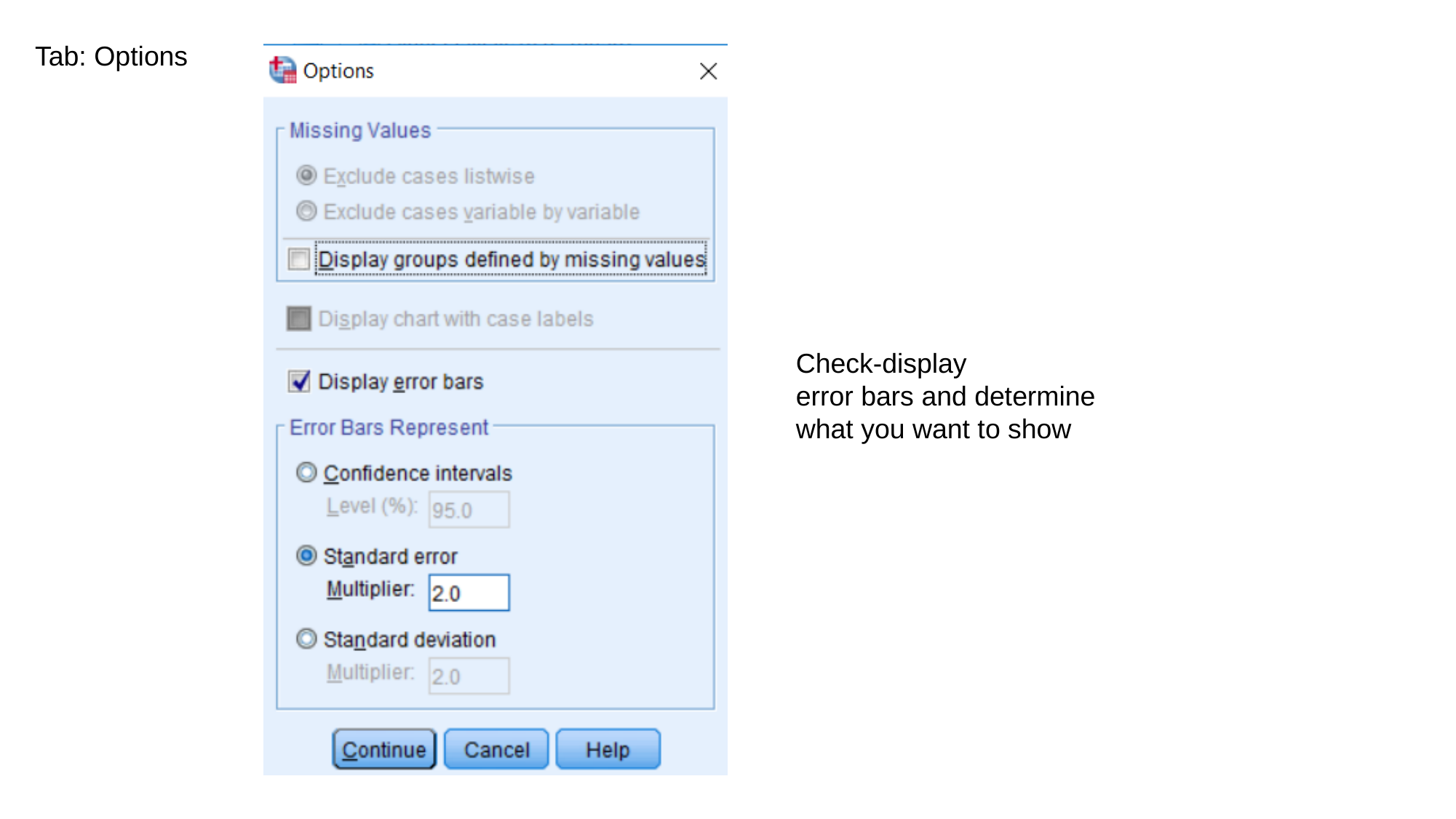

Tab: Options
Check-display
error bars and determine
what you want to show

### Slide 16
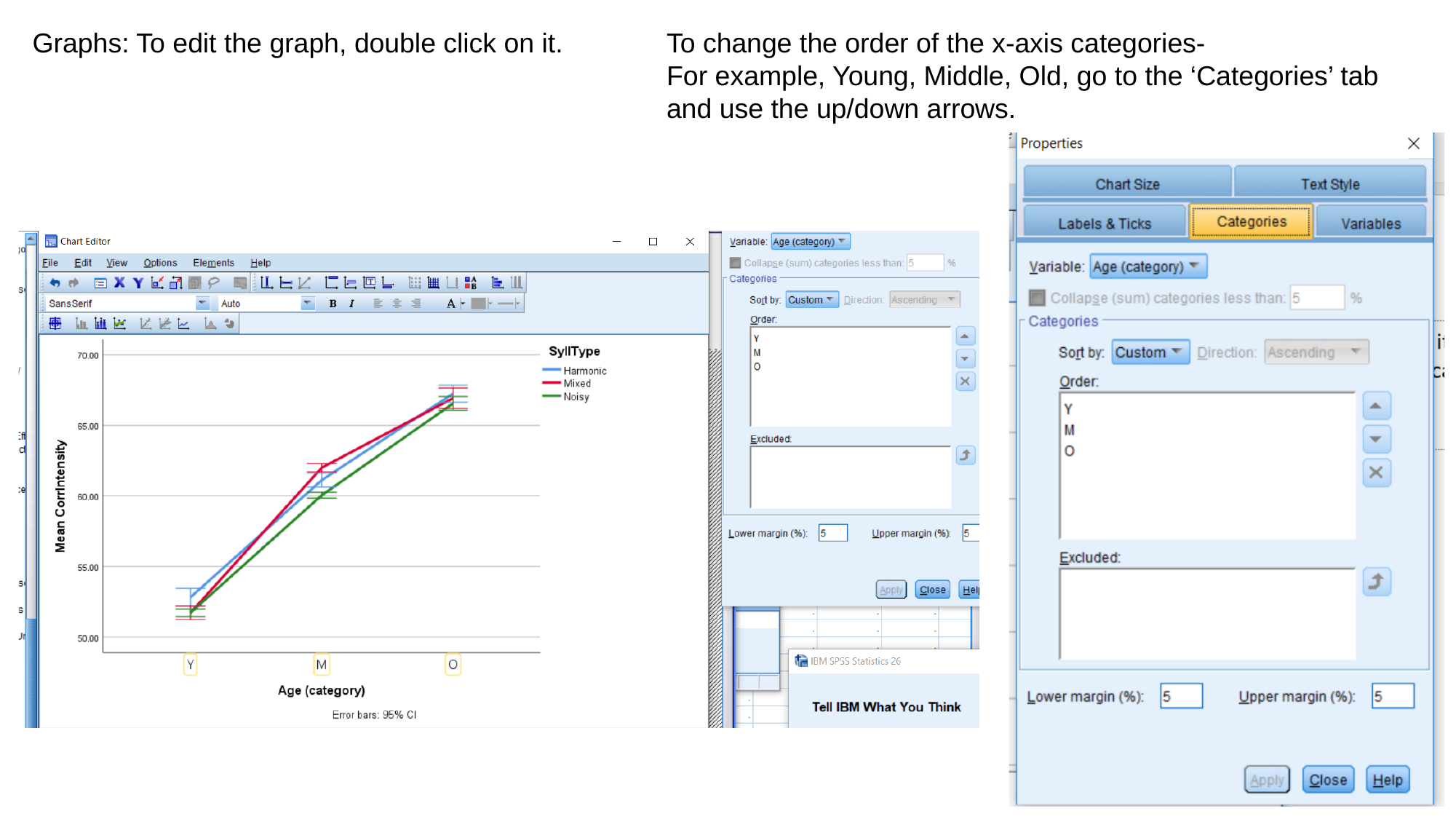

Graphs: To edit the graph, double click on it.
To change the order of the x-axis categories-
For example, Young, Middle, Old, go to the ‘Categories’ tab and use the up/down arrows.
